# 3D mapping reveals a complex and transient interstitial matrix during murine renal development

**DOI:** 10.1101/2020.08.20.258152

**Authors:** Sarah N. Lipp, Kathryn R. Jacobson, David S. Hains, Andrew L. Schwarderer, Sarah Calve

**Author notes:** Address for Correspondence: Sarah Calve, Associate Professor of Mechanical Engineering, University of Colorado - Boulder, 1111 Engineering Dr., Boulder, CO 80309.

## Abstract

**Background:** The extracellular matrix (ECM) is a network of proteins and glycosaminoglycans that provides structural and biochemical cues to cells. In the kidney, the ECM is critical for nephrogenesis; however, the dynamics of ECM composition and how it relates to 3D structure during development is unknown.

**Methods:** Using embryonic day (E)14.5, E18.5, postnatal day (P)3, and adult kidneys, we fractionated proteins based on differential solubilities, performed liquid chromatography tandem-mass spectrometry, and identified changes in ECM protein content (matrisome). Decellularized kidneys were stained for ECM proteins and imaged in 3D using confocal microscopy.

**Results:** We observed an increase in interstitial ECM that connect the stromal mesenchyme to the basement membrane (TNXB, COL6A1, COL6A2, COL6A3) between the embryo and adult, and a transient elevation of interstitial matrix proteins (COL5A2, COL12A1, COL26A1, ELN, EMID1, FBN1, LTBP4, THSD4) at perinatal timepoints. Basement membrane proteins critical for metanephric induction (FRAS1, FREM2) were highest in the embryo, whereas proteins necessary for glomerular basement membrane integrity (COL4A3, COL4A4, COL4A5, LAMB2) were more abundant in the adult. 3D visualization revealed a complex interstitial matrix that dramatically changed over development, including the perinatal formation of fibrillar structures that appear to support the medullary rays.

**Conclusion:** By correlating 3D ECM spatiotemporal organization with global protein abundance, we identified novel changes in the interstitial matrix during kidney development. This new information regarding the ECM in developing kidneys offers the potential to inform the design of regenerative scaffolds that can guide nephrogenesis *in vitro*.

**Significance statement:** End-stage renal disease is increasing and there are a limited number of organs available for transplantation. Therefore, researchers have focused on understanding how cellular signaling influences kidney development to expand strategies to rebuild a kidney. However, the extracellular matrix (ECM), another critical component that biomechanically regulates nephrogenesis, has been largely neglected. This paper combines proteomics and 3D imaging of the murine kidney to resolve previously undescribed dynamics of the interstitial matrix in the cortex and corticomedullary junction during development. Combined with cell and growth factors, scaffolds modeled after the composition and organization of the developmental ECM have the potential to improve tissue engineering models of the kidney, like organoids.

## Introduction

The increasing incidence of end-stage renal disease (ESRD) represents a worldwide public health crisis.^1^ Although the preferred treatment of ESRD is transplant, there are long wait times, indicating a critical need for additional organs.^1^ Tissue engineers aim to supplement the transplant pool by designing artificial kidneys that use polymer networks to provide an external scaffold for renal cells.^2^ Current efforts have led to incomplete differentiation *in vitro*.^3^ Therefore engineered networks may not provide the same environmental cues as the extracellular matrix (ECM), the native scaffold that integrates disparate cells into a functional kidney.

The ECM is a network of proteins and glycosaminoglycans that influences cellular behavior via biological and mechanical signaling in developing, homeostatic, and diseased tissues.^4^ ECM in the kidney can be divided into the interstitial matrix and basement membrane.^5^ The interstitial matrix is a loose network comprised of fibril-forming ECM, which includes elastin, fibrillins, collagens, proteoglycans, and glycoproteins, and is involved in maintaining mechanical integrity and growth factor sequestration.^5^ The basement membrane, a dense meshwork of macromolecules that directly surrounds cells, is critical for renal development and physiology.^4,6^ Due to the complexity of these networks, little is known how the interstitial and basement membrane matrices are assembled during nephrogenesis, generating a roadblock in implementing the appropriate microstructure and biochemical properties of the native ECM into bioengineered scaffolds.^2,7^

Traditional analyses of the ECM, (*e*.*g*. gene expression, western blotting, immunohistochemistry, knockout models, human mutation case studies), only focus on a few proteins within each study. A more comprehensive analysis to investigate the composition of complex ECM protein networks can be achieved by proteomics using liquid chromatography-tandem mass spectrometry (LC-MS/MS).^5^ Recently, fractionation techniques were developed to isolate ECM proteins, termed the matrisome, using increasingly stringent solutions to solubilize cellular proteins and characterize the matrix in diverse tissues,^8^ including fetal and adult kidneys.^9- 20^ Despite these reports, there has yet to be a study that interrogates ECM dynamics across murine embryonic, fetal, and adult kidney development.

In addition to identifying ECM proteins present in the kidney, it is critical to know how these proteins are spatially distributed. Correlating protein abundance changes with nephron structure in 3D can provide insight into potential roles for different ECM proteins. The recent development of techniques to visualize the ECM of intact tissues has provided information not apparent from 2D sections for visceral organs.^21-23^ While the 3D ECM structure of the basement membrane and developing blood vessels have been studied in the kidney,^24-26^ no reports focus on the interstitial matrix at the nephron level due to the lack of techniques to visualize deep within the tissue. We previously demonstrated that coupling immunofluorescent staining and confocal imaging with decellularization provided new insight into ECM patterning during development of the whole embryo and forelimb;^27^ however, this technique has not been applied to the kidney. In this study, we combined tissue fractionation, LC-MS/MS and 3D imaging to investigate the ECM in kidneys from the embryo to the homeostatic adult and found that the composition and organization of the interstitial matrix dramatically changed during development.

## Materials and Methods

Wild-type (WT) C57BL/6 mice were obtained from the Jackson Laboratory. Experimental protocols complied with, and were approved by, the Purdue Animal Care and Use Committee (PACUC; protocol# 1209000723). PACUC assesses that Purdue University researchers and all procedures and facilities are compliant with regulations of the United States Department of Agriculture, United States Public Health Service, Animal Welfare Act, and Purdue’s Animal Welfare Assurance. E14.5-adult kidneys were collected as described in the Supplemental Methods.

### Proteomics

#### Tissue Fractionation

Kidneys were isolated from E14.5, E18.5, P3, and male adult mice (P56). To obtain enough protein for analysis, kidneys from multiple animals and/or litters were pooled for the earlier timepoints (Table S1). Both male and female embryos and pups were used. Proteins from the different cellular compartments were isolated using the protocol developed by the Hynes lab and modified for embryonic tissue using varying detergent and ionic strengths (detailed protocol provided in the Supplemental Methods) and analyzed using MaxQuant following.^28-30^ E14.5 kidneys proteins were compared to E14.5 whole embryo data from (Jacobson *et al*., submitted).^31^

#### LC-MS/MS analysis

Protein groups were filtered using Microsoft Excel (2019) to exclude proteins identified by less than two unique and razor peptides, reverse hits, or identified as potential contaminants (with the exception of keratins, THBS1, LUM, TPM2, PFN1, and COMP). Intensities for proteins identified in only one of the three biological replicates were removed. Proteins were annotated based on the matrisome category and cellular compartment classifications (Table S1).^29,32^ Functional characterization of matrisome proteins as “basement membrane”, “interstitial matrix”, and “other” was based on^5,13,33^ with additional re-annotations based on review of histology in the literature for proteins observed in this study (Table S1). Proteins were indicated as part of the *elastin-microfibril axis* based on literature studies (Figure S1).

The percent of the matrisome identified in the IN fraction varied over kidney development and between kidney and whole embryo at E14.5. To enable comparison between timepoints and with E14.5 embryos, “scaled LFQ” values were used (*Jacobson et al*. submitted^31^; Equation 1), where *i* = protein.

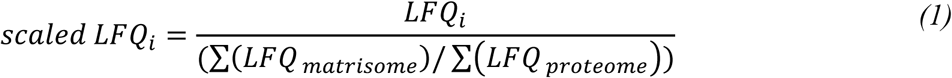

GraphPad Prism (V8.4.2), was used to visualize the data, including manually clustered heatmaps based on scaled intensity or z-score (Equation 2), where *i* = protein, *j* = sample, *S* = standard deviation.

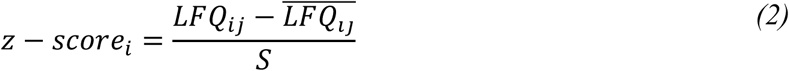

GO analysis of proteins significantly elevated or exclusive for the IN fraction for the E14.5 kidney versus whole embryo (Figure 2; Table S2) or over the kidney time course (Figure 5 and Table S1) were analyzed using g:profiler.^34^ The *Mus musculus* and *Homo sapiens* databases were used to ensure complete GO analysis annotations (*e*.*g*. laminin 11 was not found in the *Mus musculus* database), using a setting of *p* < 0.05. Groupings were modified and redundant terms were removed using REVIGO using the following setting similarity: medium (0.7), whole UniProt.^35^ The Venn diagram and figures were compiled using Adobe Illustrator.^35^

**Figure 1:**
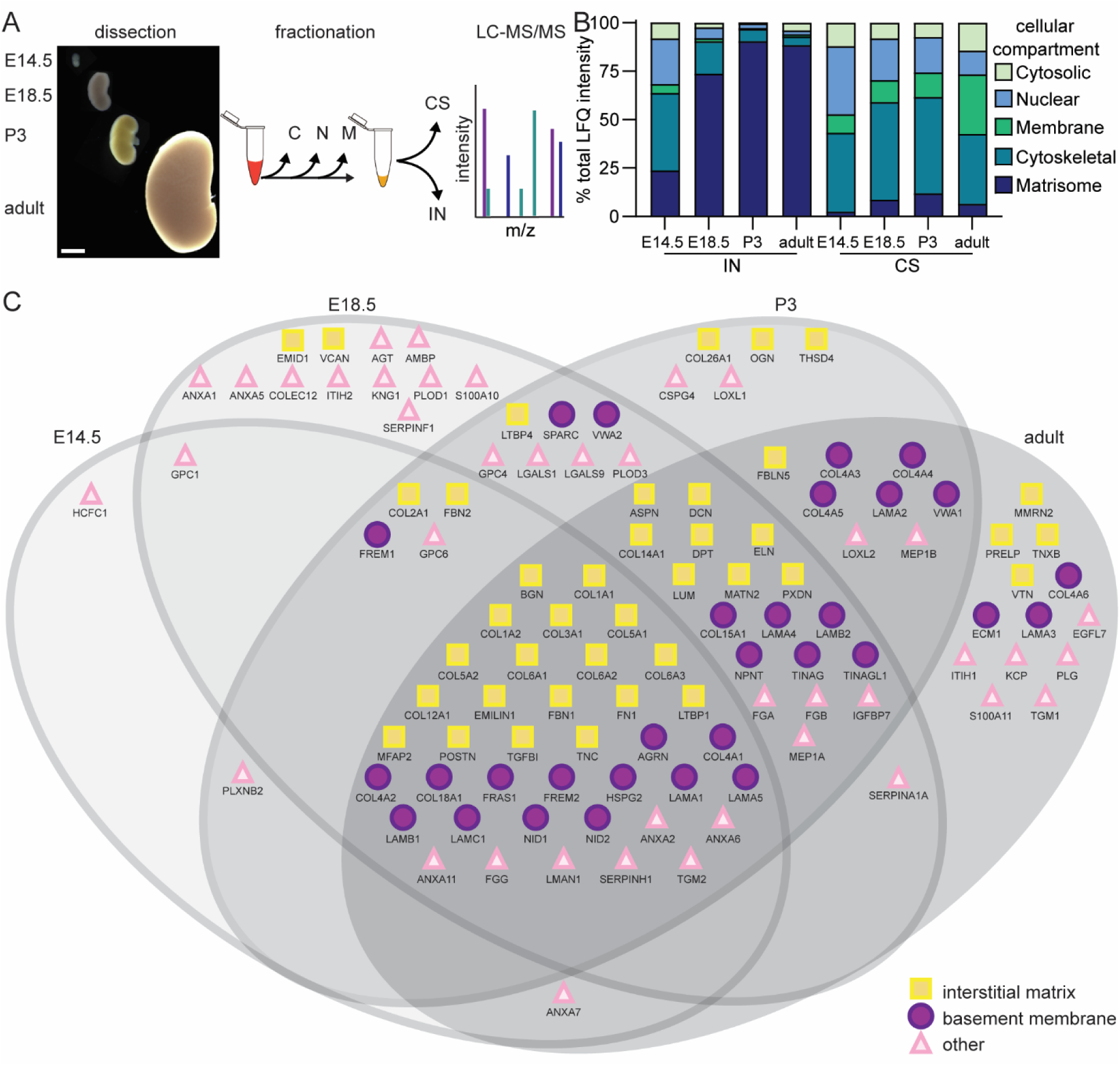
The matrisome of the murine kidney dynamically changed during development. **(A)** Embryonic day (E)14.5, E18.5, postnatal day (P)3, and adult kidneys were sequentially fractionated to isolate the insoluble (IN) and cytoskeletal (CS) fractions and analyzed using LC-MS/MS. Scale bar = 200 µm. **(B)** Comparison of total LFQ intensity of different cellular compartments showed the percent matrisome significantly varied based on developmental stage and fraction analyzed. Two-way ANOVA indicated developmental stage, fraction, and interaction were significant (*p* < 0.0001); Tukey multiple comparisons for IN fraction were significant for all comparisons (*p* < 0.0001) except for P3 versus adult. **(C)** ECM proteins found in CS and IN fractions varied between E14.5 – adult kidneys, *n* = 3 biological replicates per timepoint.

**Figure 2:**
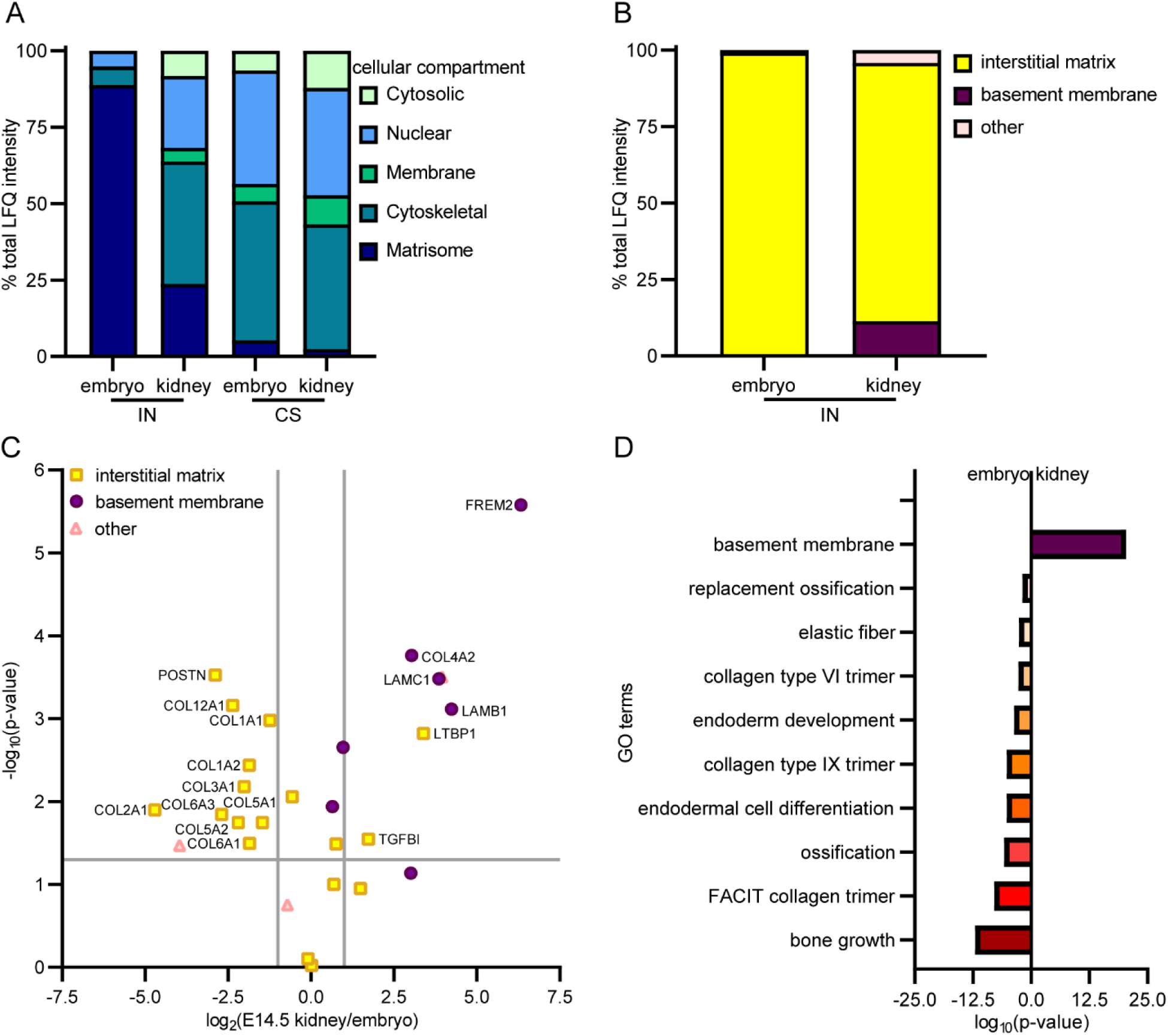
The kidney matrisome was distinct from the whole embryo by E14.5. **(A)** Comparison of total LFQ intensity of different cellular compartments showed the percent matrisome significantly varied based on tissue type and fraction analyzed. Two-way ANOVA indicated tissue, fraction, and interaction were significant (*p* < 0.0001); Sidak’s multiple comparison for IN fraction was significant (*p* < 0.0001). **(B)** There was a significant decrease in interstitial matrix and increase in the basement membrane (t-test *p* < 0.0001 for both) in the E14.5 kidney compared to the embryo. **(C)** Volcano plot comparison of log2 scaled LFQ values for IN fraction of the E14.5 whole embryo and kidney. Significance was based on *p* < 0.05 and |fold change| > 2 (grey lines). **(D)** GO analysis terms generated using proteins significantly elevated or exclusively found in the IN fractions indicate the matrisome of the kidney was specialized by E14.5 when compared with the whole embryo (*n* = 3 biological replicates per timepoint).

**Figure 3:**
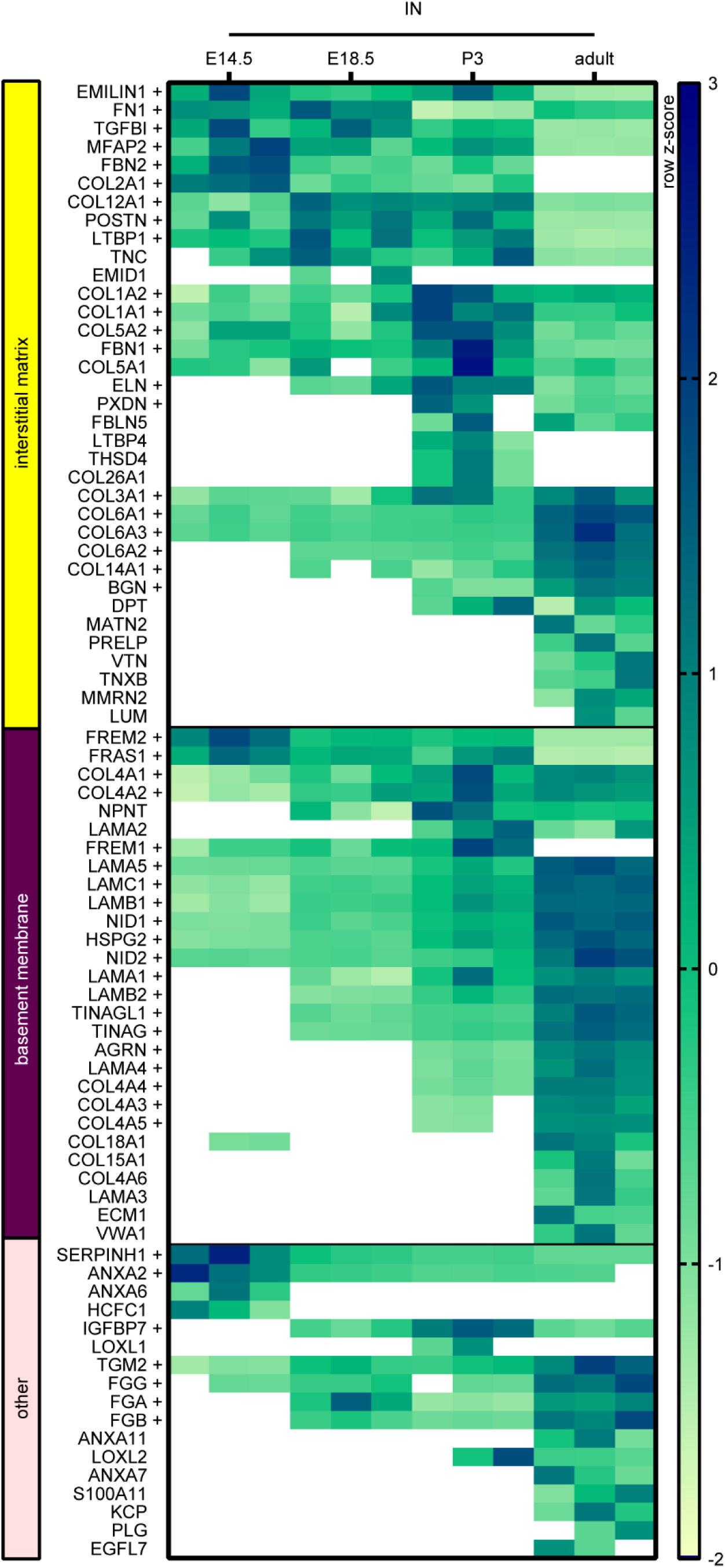
The composition of murine kidney interstitial matrix and basement membrane proteins significantly changed with development. Row z-score heat map of matrisome components, based on scaled LFQ intensity, manually grouped as interstitial matrix, basement membrane, and other ECM associated proteins for the IN fraction. + indicates *p* < 0.05 based on one-way ANOVA (identified in 3 - 4 timepoints) or unpaired, two-tailed t-test (identified in 2 timepoints). Proteins identified in *n* ≥ 2 biological replicates were included in the heat map analysis. White boxes signify zero intensity values.

**Figure 4:**
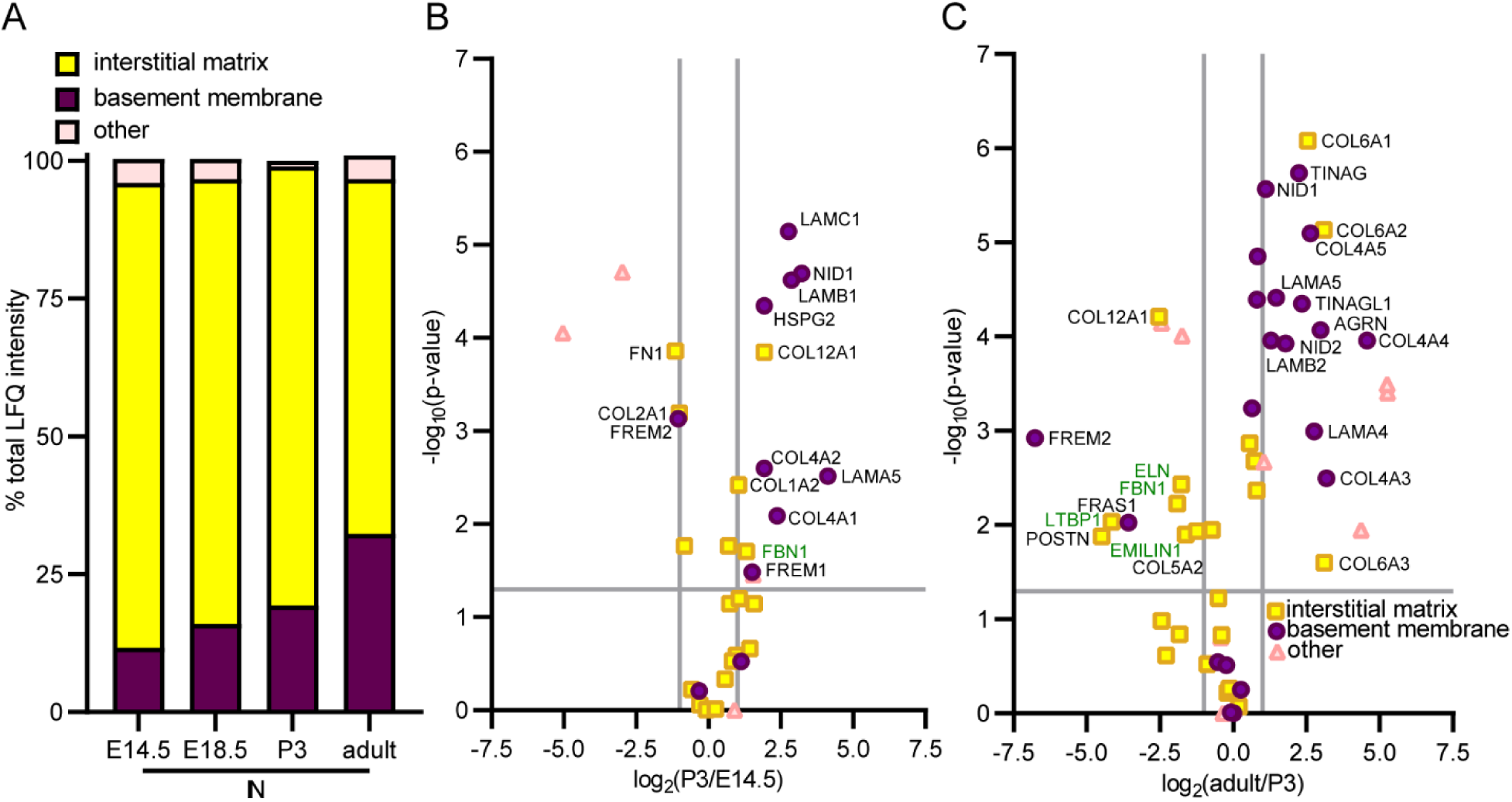
The relative composition of murine kidney interstitial matrix and basement membrane proteins significantly changed with development. **(A)** There was a significant decrease in interstitial matrix proteins and increase in basement membrane identified in the IN fraction over development. One-way ANOVA (*p* < 0.0001) and Tukey test between E14.5 and adult (*p* < 0.0001) were significant for both interstitial matrix and basement membrane. **(B, C)** Volcano plots comparing proteins found in **(B)** P3 and E14.5 or **(C)** adult and P3 IN fractions. Green protein names indicate *elastin-microfibril axis* proteins. Significance was based on *p* < 0.05 and |fold change| > 2 (grey lines; *n* = 3 biological replicates per timepoint).

**Figure 5:**
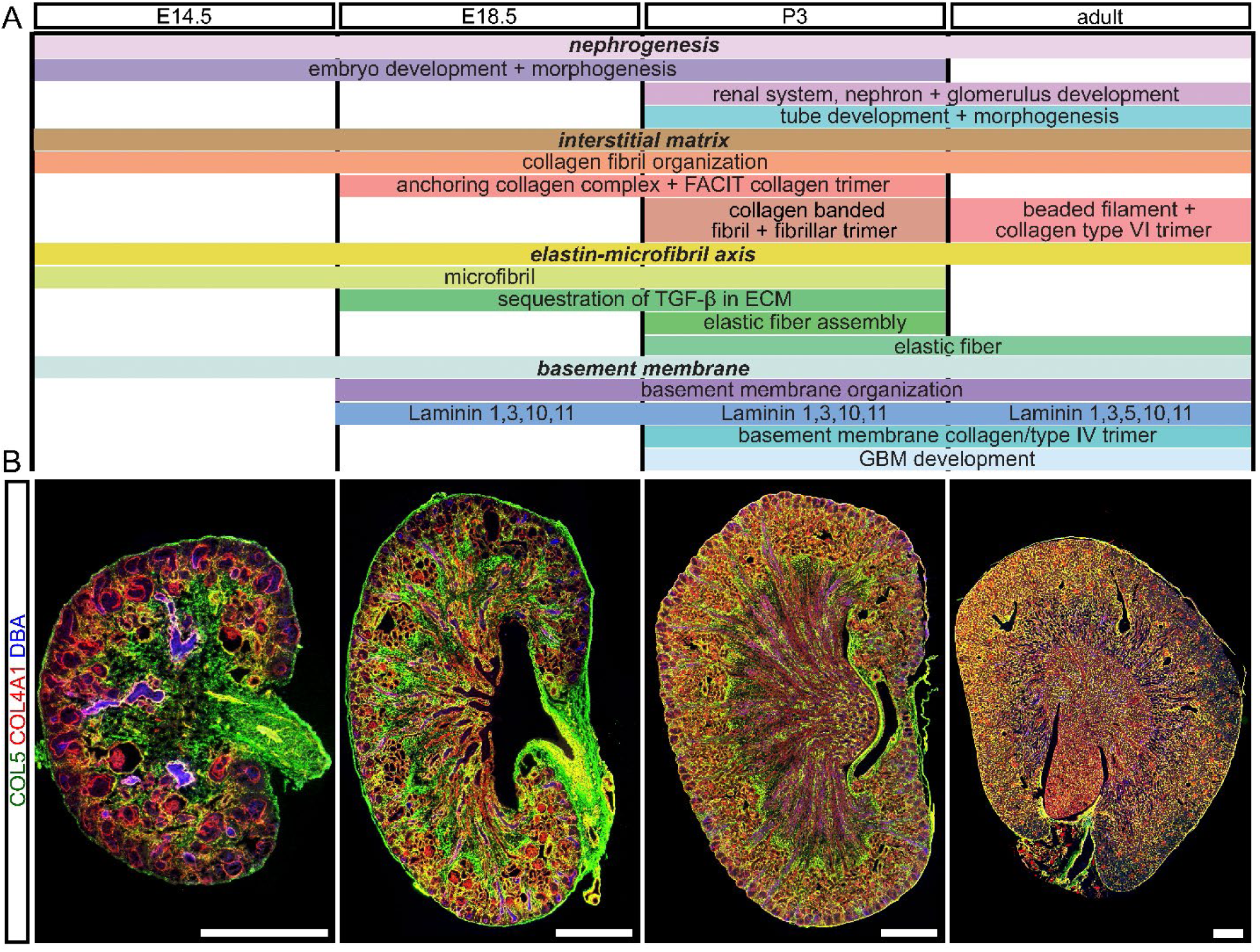
Dynamic changes in ECM function identified via GO analysis correlated with IHC. **(A)** GO analysis of proteins exclusive to, or significantly elevated in, the IN fraction (Table S1) generated “biological processes” and “cellular compartment” terms that varied over development grouped into *nephrogenesis, interstitial matrix, elastin – microfibrils axis* and *basement membrane* terms (*p* < 0.05). FACIT = fibril associated collagens with interrupted triple helices; GBM = glomerular basement membrane. **(B)** Cryosections of E14.5 – adult kidneys showed a decrease interstitial matrix space (green = COL5) and a corresponding increase in basement membrane area (red = COL4A1; blue = DBA, a collecting duct marker). Scale bars = 500 μm. Representative images from *n* = 2 biological replicates. Individual channels can be found in Figure S5.

The matrisome components identified in this study were compared with previously published studies analyzing the adult kidney,^9-12^ fetal kidney,^12^ adult glomerular tissue,^13-17^ renal artery,^18^ and kidney culture models,^36,37^ when a list of matrisome proteins identified was available using a combined human and mouse matrisome list (Table S3).^32^

### Imaging

*Immunohistochemistry (IHC)* was performed using standard methods described in the Supplemental Methods. Antibodies source and dilutions are described in Table S4.

#### 3D imaging of decellularized kidneys

Kidneys were decellularized in low concentration SDS and imaged using 3D confocal microscopy using protocols modified from^27^ (detailed methods are described in the Supplemental Methods). Antibodies source and dilutions are described in Table S4

### Statistical analyses

Kidney and whole embryo data were collected with n = 3 biological replicates and analyzed using Prism (GraphPad, V8.4.2). The effect of tissue and developmental time were studied using t-tests or ANOVA with subsequent Tukey or Sidak’s multiple comparisons test or t-test. A detailed description is provided in the Supplemental Methods.

## Results

### Proteomic analysis showed dynamic changes in matrisome composition of the developing murine kidney

To resolve ECM changes during nephrogenesis, embryonic day (E)14.5, E18.5, postnatal day (P)3, and adult mouse kidneys were sequentially fractionated and analyzed using LC-MS/MS (Figure 1A).^28,29^ Proteins in the cytoskeletal (CS) and insoluble (IN) fractions were identified using label free quantification (LFQ) in MaxQuant to allow for comparisons between samples.^30,38^ The matrisome was significantly enriched in the IN fraction and varied based on developmental timepoint (Figures 1B, S2). Of the 110 ECM proteins identified, 35% were found at all stages, of which 18 were interstitial matrix proteins and 13 were basement membrane (Figure 1C). As 63/69 of the interstitial matrix and basement membrane proteins identified were either exclusively found in the IN fraction or were at greater intensity compared to the CS (Table S1, Figure S3), the IN fraction was the focus for subsequent analyses.

### The matrisome of the kidney was distinct from the whole embryo at E14.5

To assess if the kidney matrisome is specialized by E14.5, ECM protein abundance was compared to whole embryos (Figure 2A; Table S2).^31^ There was a significantly greater amount of basement membrane proteins in the kidney (Figure 2B-C). Gene ontology (GO) analysis for proteins exclusive or significantly elevated in the kidney generated terms associated with basement membrane (Figure 2D). In contrast, the embryo matrisome generated terms linked to musculoskeletal development, indicating that the kidney matrisome is already distinct from the whole embryo by E14.5.

### The composition of murine kidney interstitial matrix and basement membrane proteins significantly changed with development

To visualize how the relative amounts of interstitial matrix, basement membrane, and ECM associated proteins changed with development, the proteins identified in the IN fraction or IN and CS were plotted as a z-score heatmap (Figures 3, S3; Table S1). Interestingly, several interstitial matrix proteins (COL5A2, COL12A1, COL26A1, EMID1, FBN1, FBLN5, ELN, THSD4) were significantly elevated or exclusively found at the perinatal timepoints, including components of the *elastin-microfibril axis*.^39-41^

Overall, there was a significant decrease in interstitial matrix and a significant increase in basement membrane proteins over development (Figure 4A). For example, the basement membrane proteins LAMA5 and NID1 were significantly increased at P3 compared to E14.5 (circles, Figure 4B), with a further increase in additional basement membrane proteins (COL4A3, COL4A4, COL4A5, LAMB2, TINAG) when comparing the adult to P3 (Figure 4C). Notably, at P3, *elastin-microfibril axis proteins* were enriched compared with the adult (green text).

### GO analysis and IHC confirms dynamic changes in matrisome abundance

GO analysis of IN fraction proteins generated timepoint-specific terms (Figure 5A; Table S1). E14.5 kidneys were ascribed GO terms that included *embryonic development* and *microfibril*. Terms assigned to E18.5 and P3 included *microfibril, sequestration of TGFβ*, and *anchoring collagen complex. Elastic fibers* and basement membrane-related terms (*collagen type IV* and *glomerular basement membrane development*) were assigned only to the P3 and adult kidney.

To validate the proteomic trends (Figure S4), cryosections of kidneys were stained with antibodies against the α1 chain of type IV collagen (COL4A1) and all chains of type V collagen (COL5), as well as *dolichos biflorus agglutinin* (DBA), a collecting duct marker (Figures 5B, S5).^42^ LC-MS/MS analysis indicated COL5A2 significantly varied over development with a peak at P3 (Figure S4). COL5 was visualized around tubules at E14.5. At perinatal timepoints, COL5 fibers were observed running in parallel with the tubules, referred to as “vertical fibers”, and around the medullary rays, referred to as “medullary ray sheath fibers”; however, in the adult, COL5 was not as prevalent and was interspersed between the tubules. In contrast, the abundance of COL4A1 significantly increased with time along with other basement membrane components (Figure S4), and the relative area of tubule basement membrane (COL4A1), compared to interstitial space (COL5), was greater in the adult than the developmental timepoints (Figure 5B).

### 3D visualization of the developing kidney revealed intricate architecture of the interstitial matrix

We previously demonstrated that 3D visualization of the ECM during murine forelimb development identified structures that were not revealed by standard 2D IHC.^27^ Therefore, to gain insight into the 3D structure of the ECM, kidneys were decellularized, stained to identify proteins of interest and imaged using confocal microscopy (Figure 6). At E14.5, when the corticomedullary junction was forming, fibers positive for COL5 surrounded the developing nephron (COL4A1) or ran parallel to blood vessels, but did not form bundles (Figures 7A-B, S6, S7; Videos S1, S2). Fibers also circumferentially surrounded the developing ureter (Video S1). By E18.5 and P3, the interstitial matrix (COL5, COL6; Figure 7C-F; Videos S1-S3) coalesced into bundles surrounding the tubules, indicated by the staining for the basement membrane components COL4A1 and HSPG2. These fibers appeared continuous with the renal calyx, pelvis, and ureter, and subdivided the medullary rays at cortical points (Videos S1, S2).

**Figure 6:**
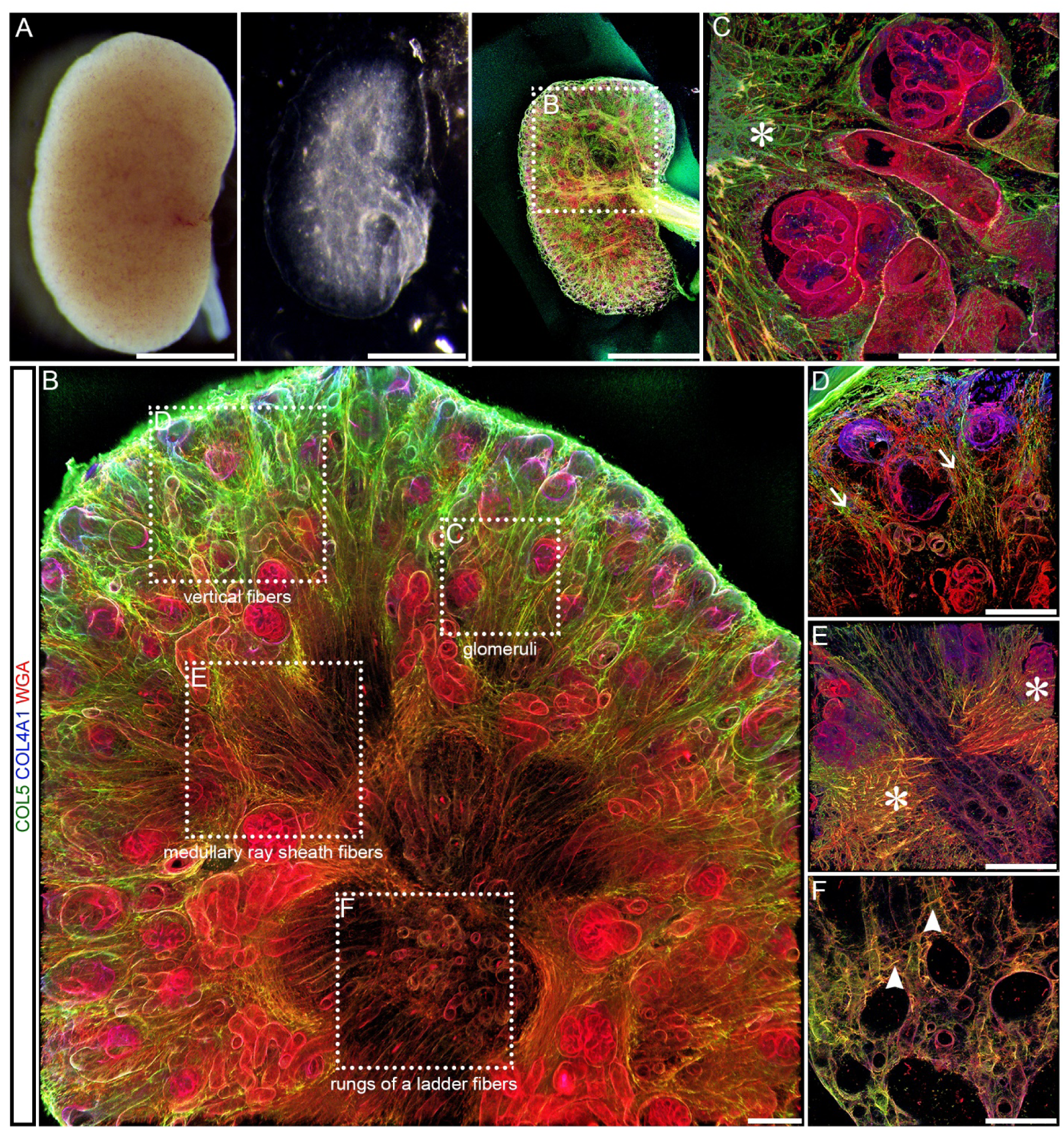
Decellularization revealed a complex 3D arrangement of the ECM during murine kidney development. **(A)** Freshly harvested kidneys (left) were decellularized in SDS (middle), then stained for ECM of interest (right; E18.5 kidney, scale bar = 1 mm). Green = COL5; blue = COL4A1; red = WGA (proteoglycans). **(B)** Fibers in the glomeruli, cortex, and corticomedullary junction could be visualized in 3D within a decellularized E18.5 kidney (10×). (**C-F)** Representative confocal images from different areas of the kidney (indicated in B with boxes) at higher magnification. **(C)** Fibrous ECM (*) surrounded glomeruli (63×). **(D)** Vertical fibers (arrow) were visualized on the cortical surface (40×). **(E)** Medullar ray sheath fibers (*) were observed at the corticomedullary junction (40×). **(F)** Fibers with a “rungs of a ladder” morphology (arrowhead) were observed in the medulla (40×). B – F: Scale bars = 100µm. Confocal z-stack dimensions in B: 1.42 mm × 1.42 mm × 100 µm (*x* × *y* × *z*); C: 225 × 225 × 37 µm; D, E: 354 × 354 × 45 µm, F: 354 × 354 × 11 µm.

**Figure 7:**
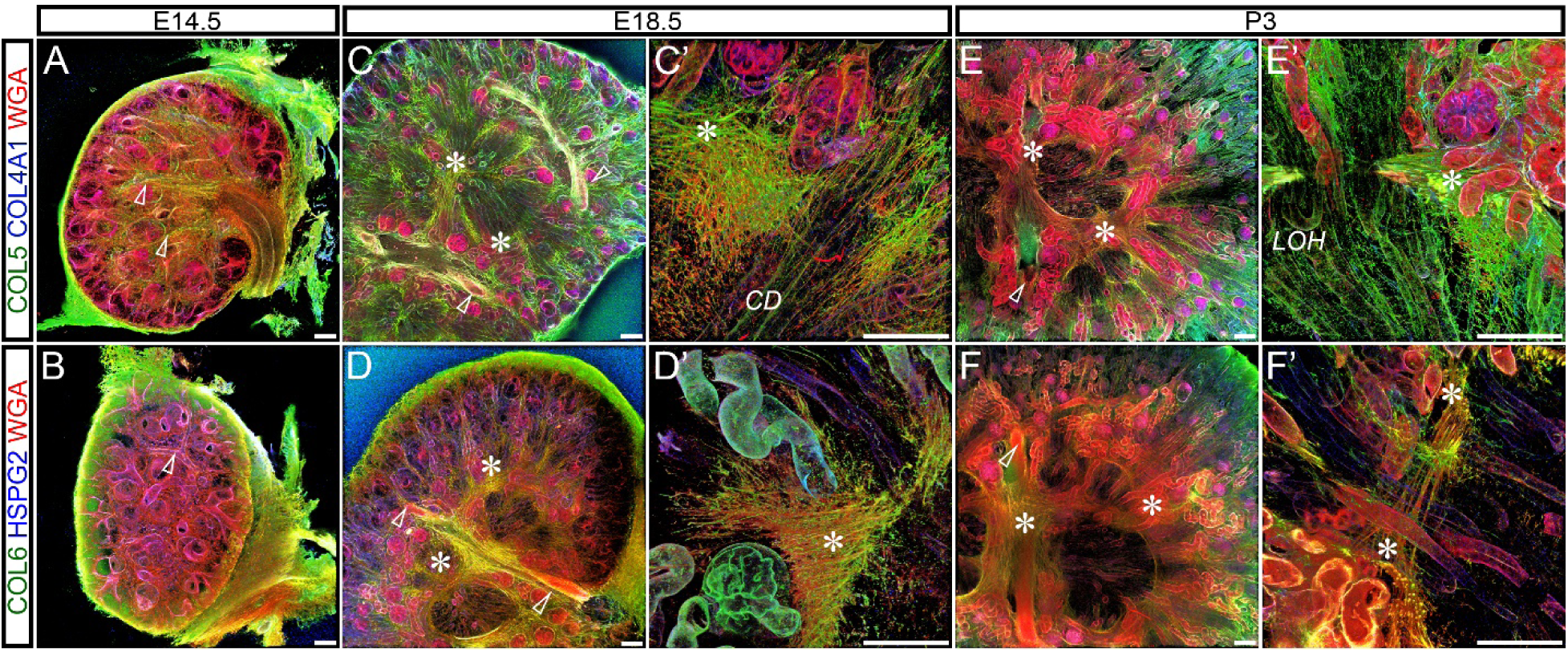
Medullary ray sheath fibers were only found at E18.5 and P3. **(A-B)** At E14.5, a fibrous ECM (green = COL5 and COL6) ran parallel to developing blood vessels and surrounding tubules (blue = COL4A1, HSPG2; red = WGA). **(C - F’)** Medullary ray sheath fibers (*) surrounded the developing nephron (*CD* = collecting duct, *LOH* = loop of Henle) at E18.5 – P3. Scale bars = 100 µm. Dimensions of 10× confocal z-stacks in A, B, C, D, E and F: 1.42 mm × 1.42 mm × 100 µm (*x* × *y* × *z*); in C’ and E’ 40× insets: 354 × 354 × 45 µm; in D’ and F’ 40× insets: 354 × 354 × 36 µm. Open arrowheads = blood vessels. Representative images from *n* = 3 biological replicates. Individual channels can be found in Figures S6, S7.

Fibers at E14.5 emanated from, and ran parallel to, the subcapsular tubules to the capsule. At E18.5 and P3, fiber bundles ran parallel (COL5) to the developing nephron and tubule (COL4A1, HSPG2). Vertical fibers were observed from the base of the medullary ray sheath fibers to the nephrogenic cortex, where the vertical fibers intertwined at the surface (Figures 8A, C, E, S9; Videos S1, S2, S4). In contrast to the cortex, fibers in the medulla were perpendicular to the tubules in a “rungs of a ladder” configuration (Figures 8B, D, F, S9, S10). In the adult, bulk accumulation of fibers or clear cortical or medullary patterning was not observed, but “rungs of a ladder” structures were detected (Figure S11). The 3D imaging revealed dynamic structures of the interstitial extracellular matrix that have not been previously described in the cortex and corticomedullary junction at perinatal timepoints.

**Figure 8:**
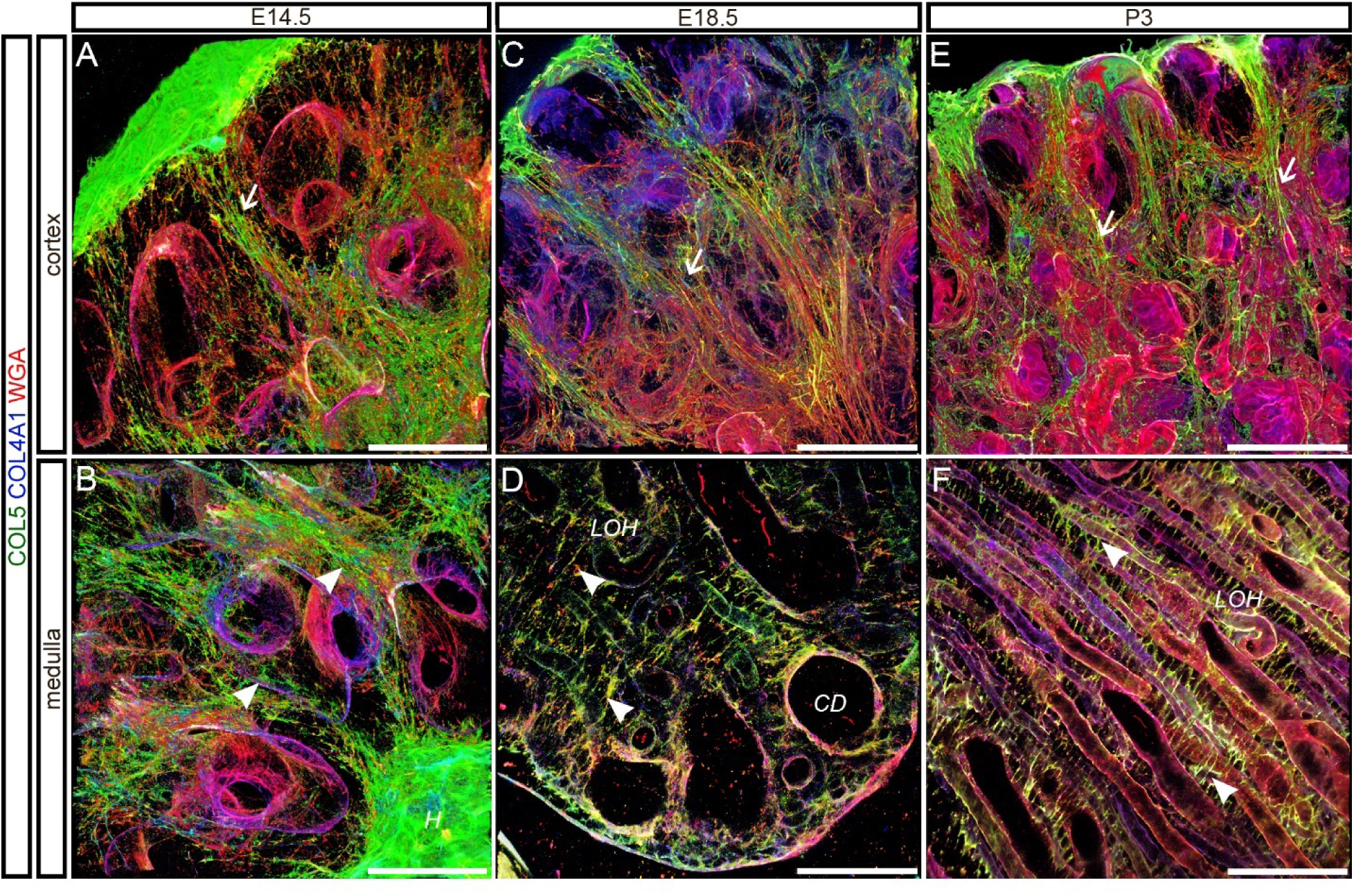
The morphology of interstitial fibers was distinct between the cortex and medulla. **(A, C, E)** E14.5, E18.5, and P3 vertical fibers (arrows) ran parallel to, and emanated from, the tubules to the capsule in vertical fibers (green = COL5; blue = COL4A1; red = WGA). (**B**) Some E14.5 fibers (arrows) extended from the developing renal hilum (*H*). **(D, F) “**Rungs of a ladder” appearance of the interstitial matrix (arrowheads) at E18.5 and P3 between loop of Henle (*LOH*) and collecting duct (*CD*) tubules. Scale bars = 100 µm. Dimensions of 40× confocal z-stacks in A, B, C, E: 354 × 354 × 45 µm (*x* × *y* × *z*); D, F: 354 × 354 × 11 µm. Representative image from *n* = 3 biological replicates. Individual channels can be found in Figures S8, S9.

## Discussion

A paucity of information about the interstitial matrix in the kidney was noted over 20 years ago.^43^ While components such as FN1 and FBN1 were shown to be critical for normal branching morphogenesis in explant culture in the intervening years,^44,45^ the role of the interstitial matrix in renal development *in vivo* is still unclear. To begin to address this gap in knowledge, we combined proteomic analyses and 3D imaging to map ECM dynamics during kidney development. We observed some interstitial matrix proteins with a role in mechanical strength increased over development, correlating with increased hydrostatic pressure,^46^ while *elastin–microfibril axis* proteins were transiently elevated (Figures S3, 4). To further interrogate the changes in ECM composition, we used 3D imaging and observed transient cortical medullary ray structures at the perinatal timepoints and vertical fibers in the cortex (Figures 7, 8; Videos S1-S4).

### The composition and structural arrangement of the interstitial matrix dynamically changed over development

Overall, we observed a global decrease in interstitial matrix proteins in the adult as determined by mass spectrometry (Figure 4). FN1, a glycoprotein with a role in orchestrating ECM assembly^47^, was enriched at early timepoints (Figure 3), correlating with the critical role in promoting branching in explant cultures.^44^ Type II collagen (COL2A1), classically associated with cartilage formation^48^, was also elevated in the developing kidney (Figure 3). COL2A1 forms homotrimers and polymerizes into fibrils that provide strength in tension. COL2A1 may play a role in supporting the developing renal epithelium by enhancing the strength of attachment to the underlying mesenchyme.^49^

In the E14.5 kidney, our 3D analysis showed type V collagen (COL5) fibers circumferentially surrounding the ureteric bud (Video S1) in a similar pattern to the layers of primary stromal mesenchyme.^50^ Type V collagen is critical for type I collagen fibrillogenesis^51^, and forms fibrils that provide resistance to tensile forces, similar to COL2A1, contributing to tissue strength.^52^ Vertically aligned fibers, containing COL5, were found in the cortex of E14.5, E18.5, and P3 kidneys running parallel to the nephron, extending perpendicular from the corticomedullary junction fibers to the capsule, and intertwining at the surface (Figures 7, 8, S6, S8; Videos S1, S4). Although we observed a few fibers connecting the collecting duct ampule to the capsule as previously described,^53^ the majority of interstitial ECM envelop the nephron elements in vertical fibers. Like the ampule fibers, we hypothesize the vertical fibers regulate tissue stability as well as maintain the radial growth of the collecting duct due to connection to the capsule. Fibers bridging the collecting duct and loop of Henle in the medulla were observed at E18.5 to adult (Figures 8, S9, S11), similar to the “rungs of a ladder” orientation of stromal cells previously described.^54^ We hypothesize the COL5 positive ECM structures are involved in establishing and maintaining the tubular arrangement of the loop of Henle, vasa recta, and collecting duct to facilitate filtrate concentration in the inner medulla.^55^

At E14.5, ECM fibers ran parallel to blood vessels, which may remodel into the interconnected rings of medullary ray sheath fibers at the perinatal timepoints located at the corticomedullary junction (Figure 7). These COL5 positive interstitial matrix structures could provide bundling structural support of the loop of Henle. The location of the medullary ray sheath fibers correlate with the organization of the primary stromal mesenchymal cells, which become restricted to areas between the medullary rays.^56-58^ Between perinatal and P28 timepoints, the space occupied by the primary mesenchymal stromal cells is thought to disappear due to large scale apoptosis or decreased proliferation and was replaced by the loop of Henle and vasa recta during the radial outgrowth of the nephron cascade.^54,56,59-61^ However, it is unclear if the medullary ray sheath fibers are remodeled to surround the nephron or are degraded.

At the same perinatal timepoints as the medullary ray sheath fibers were observed, LC-MS/MS revealed some interstitial matrix proteins were transiently increased. These proteins include COL5A2 and type XII collagen (COL12A1), which facilitates collagen organization by bridging collagen fibrils with other interstitial matrix proteins (Figure 4).^62,63^ Additional proteins that were transiently upregulated include those associated with the *elastin-microfibril axis* (COL26A1, ELN, EMID1, FBN1, LTBP4, THSD4; Figures 3, 4C). ELN provides tissue compliance and fibrillins (FBNs) form microfibrils that resist tensile loads, act as a scaffold for elastic fibril formation, and sequester growth factors (*e*.*g*. members of the TGFβ superfamily).^64^ Notably, GO analysis showed *microfibril, TGFβ sequestration*, and *elastin* terms overlapped only at the P3 timepoint (Figure 5). The perinatal increase in abundance of these collagens and *elastin-microfibril axis* proteins may provide mechanical integrity by transiently stabilizing the 3D fibrous medullary ray sheath and vertical fibers (Figures 5, 7, 8). Interestingly, proteins in the *elastin-microfibril axis* (EMILIN1, EMILIN3, FBN1, FBN2, FBN3, and THSD4)^39-41^ were also elevated in human fetal kidneys compared with the adult,^12^ indicating this understudied axis in the kidney warrants further research.^40,41^

In contrast with the fibrous structures observed at perinatal timepoints, relatively little interstitial patterning was observed in the adult using our decellularization method (Figure S11). Complex structures may be better resolved in the adult if perfusion decellularization is used instead of passive decellularization protocols since previous electron microscopy studies showed fibers near the tubules and capillaries.^24,65^ The ultrastructural organization of fibers correlates with the increase in proteins interacting with the basement membrane we observed in the adult matrisome. TNXB, a ECM glycoprotein implicated in the regulation of collagen fibril spacing,^66^ was found only in the adult (Figure 4). The increase in TNXB could provide support related to the increased hydrostatic pressure with kidney maturation,^46^ as it was suggested that TNXB was necessary for closing the ureterovesical junction during voiding.^67^ Likewise, type VI collagen, a network-forming collagen made up of COL6A1, COL6A2, and COL6A3 chains, bridges the interstitial matrix to the basement membrane,^68^ and is significantly increased in the adult (Figure 3). A patient with recurrent hematuria had a mutation in COL6A1, suggesting a disruption of COL6A1 lessened the anchoring of the interstitial matrix to, and increased proteolytic damage of, the basement membrane.^69^ This indicated that COL6A1 could be required to withstand the increased pressure found in the mature kidney. Small leucine-rich proteoglycans (ASPN, BGN, DCN, LUM, PRELP, OGN), proteins that sequester growth factors and connect within the interstitial matrix and to the basement membrane, also increased in adult tissue (Figure S3).^70,71^ The increase in these proteoglycans could be a physiological response to the increased hydrostatic pressures observed in the adult.^46^ This supposition is based on the role of small leucine-rich proteoglycans in providing strength in other tissues^70^ and BGN knockout mice exposed to increased hydrostatic pressure had cystic dilation of tubules and Bowman’s capsule potentially resulting from weak connective tissue.^72-74^

### The basement membrane dynamically changed in composition with renal development

A comparison of the ECM found in whole embryos and kidneys showed that by E14.5, the kidney had significantly more basement membrane components and fewer proteins associated with skeletal development (Figure 2). Fraser syndrome complex proteins (FRAS1, FREM2) were elevated in the developing kidney relative to the whole embryo and adult kidney (Figures 2, 3, 4). The FRAS complex promotes epithelial-mesenchymal interaction during metanephric induction with mutations causing renal agenesis.^75,76^ The early differentiation of the basement membrane can explain the strong renal phenotype for patients with FRAS complex protein mutations.^77^

Overall, the abundance and complexity of the basement membrane matrisome increased in the adult kidney compared to developmental timepoints (Figures 3, 4, 5). The general increase in basement membrane proteins (AGRN, COL4A1, COL4A2, HSPG2, LAMA1, LAMA4, LAMB1, LAMC1, NID1, NID2, TINAG, and TINAGL1) is likely due to the increase in total basement membrane volume as shown in 3D for COL4A1 and HSPG2 (Figures 5, 7, S6, S7; Videos S1, S2), as well as the previously described increase in thickness with age.^78-80^ This change in geometry could be a physiological response to increasing hydrostatic pressure and urine output over the course of development.^46,81^ COL4A1 and HSPG2 are necessary for basement membrane integrity as mutations can lead to tubular basement membrane splitting and cysts, glomerulocystic disease, and altered GBM function.^82-84^ Proteins known to promote glomerular filtration unit stability and glomerular maturation,^85-89^ COL4A3, COL4A4, COL4A5, LAMA3, LAMA5, and LAMB2, were significantly elevated or exclusive to the adult (Figures 3, 4C). The increase correlated to the incorporation of these chains into the mature GBM collagen and laminin trimers during the capillary loop stage of glomerulogenesis.^88,90^ Mutations in *COL4A3-5* can yield Alport syndrome that manifests as glomerular hematuria,^86^ while mutations in *LAMB2* and *LAMA5* yield nephrotic syndrome.^87,89^

### A common kidney matrisome was observed across multiple proteomic studies

The ECM proteins we identified had a large percentage of overlap with previous proteomic studies;^9- 18,36,37^ however, there were some differences (Table S3). Variations could be due to different tissue fractionation methods, protein abundance, and sample type. For example, our study did not resolve COL6A4-6, DMBT1, and FBLN1 (Table S3).^91-93^ These proteins were potentially extracted in the less stringent buffers at earlier timepoints as the ECM was still being incorporated into a cohesive network.^31^ This is supported by the observation that some enzymes involved in ECM crosslinking,^4,94^ including LOXL1, LOXL2, PXDN, TGM1, and TGM2, increased in abundance in the postnatal and adult kidney (Figure S3; Table S1).

## Conclusion

By combining proteomics and 3D imaging of decellularized kidneys, we were able to resolve previously undescribed dynamics of the interstitial matrix. We demonstrated that the kidney matrisome is already distinct from the whole embryo by E14.5. We also identified the transient presence of vertical and medullary ray sheath fibers in the perinatal kidney. Based on the 3D ECM patterning and proteomic data, we generated a map of the ECM during renal development (Figure 9). Outstanding questions remain regarding how interstitial matrix proteins regulate kidney morphogenesis through structural support and growth factor signaling. Future studies on the role of the interstitial matrix will provide additional insight into renal development and improve the design of *in vitro* models of nephrogenesis.

**Figure 9:**
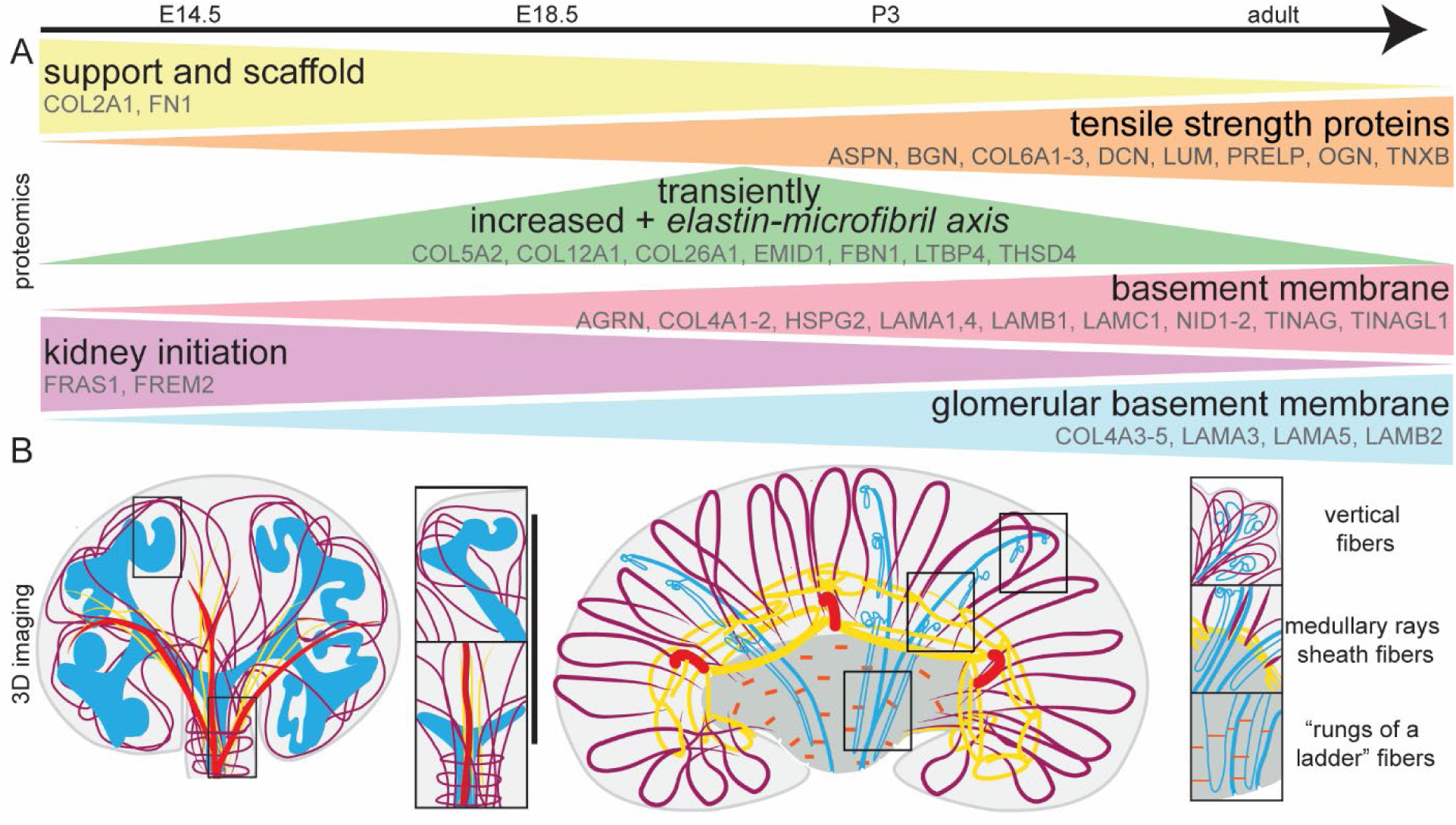
Summary of the proteomic trends and 3D ECM distribution observed. **(A)** Interstitial matrix and basement membrane proteins changed dynamically with development for proteins involved in support and scaffold, tensile strength and small leucine rich proteoglycans, transiently upregulated and *elastin-microfibril axis*, kidney initiation, basement membrane, and glomerular basement membrane. **(B)** 3D interstitial matrix was patterned at E14.5 (left) with fibers (purple) circumferentially surrounding the ureteric bud (cyan), extending parallel (yellow) to the blood vessels (red), and surrounding the developing nephron (cyan) and connecting to the capsule. At the perinatal timepoints (right), vertical fibers (purple) interconnected at the cortical surface and ran to the corticomedullary junction. At the corticomedullary junction, medullary ray sheath fibers (yellow), surrounded the developing nephron (cyan). In the medulla, fibers in a “rungs of a ladder” orientation were observed (orange).

## Data availability statement

The data supporting the findings of this study are openly available in the MassIVE repository at MSV000085616. The whole embryo E14.5 data were derived from (Jacobson *et al*., submitted) and is included in the MassIVE upload.^31^

## Acknowledgements

Ms. Lipp and Dr. Calve designed the experiments with input from Ms. Jacobson, Dr. Hains, and Dr. Schwaderer. Ms. Lipp performed the experiments; Ms. Lipp and Dr. Calve analyzed and interpreted the data and wrote the manuscript with input from Ms. Jacobson, Dr. Hains, and Dr. Schwaderer.

The authors would like to thank the Purdue Proteomics Facility and members of the Calve lab for helpful discussions, assistance with proteomics (Alexander Ocken and Naagarajan Narayanan), and confocal microscopy (Yue Leng and Andrea Acuña). In addition, the authors thank Jennifer Anderson for help with the development of the perfusion protocol.

## Supplemental Material

Supplemental Appendix 1: Supplemental Methods

Video S1: 10× medullary ray sheath fibers at E14.5, E18.5, and P3 stained for COL5, COL4A1, and WGA

Video S2: 10× medullary ray sheath fibers at E14.5, E18.5 and P3 stained for COL6, HSPG2, and WGA.

Video S3: 40× medullary ray sheath fibers at E18.5 and P3 stained for COL5, COL4A1, and WGA.

Video S4: 40× vertical fibers in the cortex of E14.5, E18.5, and P3 kidney stained for COL5, COL4A1, and WGA.

Table S1: Raw data supporting the comparison of the matrisome during kidney development.

Table S2: Raw data supporting the comparison of the matrisome between E14.5 kidneys and whole embryos.

Table S3: A common kidney matrisome was observed across different studies.

Table S4: Materials and settings used for imaging the kidney ECM.

Figure S1: Flow diagram of the functional classification of proteins as interstitial matrix, basement membrane, and other ECM associated proteins for matrisome identified in this study.

Figure S2: Proteomic techniques showed high reproducibility between the matrisome for biological replicates.

Figure S3: Heat map comparison of the dynamic changes of the matrisome for both the IN and CS fraction.

Figure S4: Scaled LFQ values for the IN fraction for proteins of interest visualized using IHC. Figure S5: Spatiotemporal change of COL5, COL4A1 in murine developing kidney.

Figure S6: Medullary ray sheath fibers were only found at E18.5 and P3 and stained for COL5.

Figure S7: Medullary ray sheath fibers were only found at E18.5 and P3 and stained for COL6.

Figure S8: COL5 vertical fibers were observed in the cortex.

Figure S9: COL5 “rungs of a ladder” fibers were observed in the medulla.

Figure S10: The morphology of interstitial fibers was distinct between cortex and medulla for COL6.

Figure S11: Interstitial accumulations were not observed in the adult mouse kidney.

