## Supplementary material for "3D mapping reveals a complex and transient interstitial matrix during murine renal development": methods and supplemental figures

**Table of contents for supplemental material**


Supplemental Appendix 1: Supplemental Methods

Video S1: 10× medullary ray sheath fibers at E14.5, E18.5, and P3 stained for COL5, COL4A1, and WGA

Video S2: 10× medullary ray sheath fibers at E14.5, E18.5 and P3 stained for COL6, HSPG2, and WGA.

Video S3: 40× medullary ray sheath fibers at E18.5 and P3 stained for COL5, COL4A1, and WGA.

Video S4: 40× vertical fibers in the cortex of E14.5, E18.5, and P3 kidney stained for COL5, COL4A1, and WGA.

Table S1: Raw data supporting the comparison of the matrisome during kidney development.

Table S2: Raw data supporting the comparison of the matrisome between E14.5 kidneys and whole embryos.

Table S3: A common kidney matrisome was observed across different studies.

Table S4: Materials and settings used for imaging the kidney ECM.

Figure S1: Flow diagram of the functional classification of proteins as interstitial matrix, basement membrane, and other ECM associated proteins for matrisome identified in this study.

Figure S2: Proteomic techniques showed high reproducibility between the matrisome for biological replicates.

Figure S3: Heat map comparison of the dynamic changes of the matrisome for both the IN and CS fraction.

Figure S4: Scaled LFQ values for the IN fraction for proteins of interest visualized using IHC.

Figure S5: Spatiotemporal change of COL5, COL4A1 in murine developing kidney.

Figure S6: Medullary ray sheath fibers were only found at E18.5 and P3 and stained for COL5.

Figure S7: Medullary ray sheath fibers were only found at E18.5 and P3 and stained for COL6.

Figure S8: COL5 vertical fibers were observed in the cortex.

Figure S9: COL5 "rungs of a ladder" fibers were observed in the medulla.

Figure S10: The morphology of interstitial fibers was distinct between cortex and medulla for COL6.

Figure S11: Interstitial accumulations were not observed in the adult mouse kidney.

**Supplemental Appendix 1:** Supplemental Methods

**Tissue collection.**

WT C57BL/6J mice were time mated, where embryonic day (E)0.5 refers to noon of the day when the copulation plug was noted. Adult mice and pregnant dams (E14.5, E18.5) were euthanized by $CO_2$ inhalation and confirmed via cervical dislocation. E18.5 and P3 pups were euthanized using decapitation. For proteomic analysis, P3 and adult mice were perfused with 1× PBS using a cardiac approach to remove blood. Embryos, P3 pups, and adult kidneys were transferred to chilled PBS on ice for fine dissection of the kidney. The ureter, capsule, and renal vein and artery were removed from the kidneys at all timepoints. Tissues to be analyzed for LC-MS/MS were snap-frozen in liquid nitrogen and stored at -80°C until processed. Kidneys for decellularization were processed immediately after harvest (as described below), and those for cryosectioning were embedded in optimal cutting temperature compound (OCT; Electron Microscopy Sciences) and stored at -80°C until cryosections were collected.

**Proteomics.**

*Proteomic isolation method:* The proteome from the different cellular compartments were isolated using the protocol developed by the Hynes lab and modified for embryonic tissue using a Compartment Protein Extraction Kit (Millipore EMD).[1,2] Halt Protease Inhibitor Cocktail (1×, Thermo Scientific) was added to all buffers and PBS and Benzonase/Nuclease B (EMD Millipore, at 1:1000) was added to the nuclear and membrane buffers. The samples were suspended in ice-chilled cytosolic buffer (C) at a ratio of 9.5:1 buffer (μL) to wet weight (mg) and homogenized (TissueRuptor, Qiagen). Samples were nutated for 30 minutes at 4°C, 400 μL of the homogenate was then centrifuged for 20 minutes at 21,100 × g at 4°C. The supernatant (C fraction) was snap-frozen and stored at -80°C. The pellet was resuspended in wash (W) buffer at a buffer (μL) to wet weight (mg) ratio of 19:1, nutated at 4°C for 5 minutes, centrifuged for 20 minutes at 21,100 × *g* at 4°C, and the supernatant removed.

The pellet was resuspended in nuclear buffer (N) an at a buffer (µL) to wet weight (mg) ratio of 4.75:1, nutated at 4°C for 5 minutes, centrifuged for 20 minutes at 21,100 × *g* at 4°C, and the supernatant collected (N1), snap-frozen and stored at -80. The pellet was resuspended in nuclear buffer (N) at a buffer (µL) to wet weight (mg) ratio of 4.75:1, nutated at 4°C for 5 minutes, centrifuged for 20 minutes at 21,100 × *g* at 4°C, and the supernatant collected (N2), snap-frozen and stored at -80. The pellet was resuspended in membrane buffer (M) at a buffer (µL) to wet weight (mg) ratio of 4.75:1, nutated at 4°C for 5 minutes, centrifuged for 20 minutes at 21,100 × *g* at 4°C, and the supernatant collected, snap-frozen and stored at -80. The pellet was then resuspended in cytoskeletal (CS) buffer at a buffer (µL) to wet weight (mg) ratio of 2.38:1, nutated at (room temperature) RT for 20 minutes and centrifuged for 20 minutes at 21,100 × *g* at 4°C, and the supernatant collected (CS1) snap-frozen and stored at -80°C. The insoluble pellet was washed with the C buffer at a buffer (µL) to wet weight (mg) ratio of 7.13:1 and centrifuged for 20 minutes at 21,100 × *g* at 4°C, and the supernatant collected (CS2). The insoluble pellet was washed in PBS at a buffer (µL) to wet weight (mg) ratio of 9.5:1 and centrifuged for 20 minutes at 21,100 × *g* at 4°C, the supernatant discarded and the insoluble pellet snap-frozen and stored at -80°.

*Peptide preparation for LC-MS/MS:* The CS buffer supernatants were combined (CS1 + CS2), and protein was precipitated in a solution of methanol, chloroform, and HPLC grade water at a ratio of 8 CS solution: 6 methanol: 3 chloroform: 8 water. The mixture was vortexed then centrifuged for 5 minutes at 18,000 × *g* at RT. The aqueous and organic phases were decanted, and the pellet was dried at RT. Insoluble pellets from the combined CS fractions and insoluble fractions were resuspended in 8M urea in 100mM ammonium bicarbonate (42.1 mg starting wet weight: 50

μL urea/ammonium bicarbonate) and reduced with dithiothreitol (VWR, final concentration 10mM) for at least 2 hours at 37°C with constant agitation. After equilibrating to RT, samples were protected from light while alkylated with iodoacetamide (VWR, final concentration 25mM) for 30 minutes. Samples were diluted to 2M urea with 100mM ammonium bicarbonate and deglycosylated with chondroitinase ABC (Sigma, final concentration 0.2U/200μL for 2 hours at 37°C with constant agitation. Peptidases were added for protein digestion: Endoproteinase LysC (1μg/200μL; New England Biolabs) for 2 hours, MS-grade trypsin (3μg/200μL; ThermoFisher Scientific) overnight, MS-grade trypsin (1.5μg/200μL) for 2 hours all conducted at 37°C with constant agitation. Trifluoroacetic acid (final concentration 0.1%) was added to inactivate digestion enzymes.

*Removal of peptide contaminates and sample preparation:* Detergent contamination was removed using 0.5 mL Pierce Detergent Removal Spin Columns (Thermo Fisher Scientific) and a modified manufacturer's protocol: After removal of the storage solution by centrifugation at 1000 × g for 1 min, the columns were prepared by washing with 400 mL of PBS and centrifuging at 1000 × g for 1 min, repeated 3 times. The digested sample was added to the column, incubated for 2 min, and centrifuged at 1000 × g for 2 min. Salts were removed with C-18 MicroSpin columns (The NestGroup, Inc.). Columns were prepared with 1 wash of 100 μL of 100% acetonitrile and 2 washes of 100 μL 0.1% trifluoroacetic acid in HPLC-grade water with centrifugation at 500 × g after each wash. Samples were loaded and spun for 8 minutes at 500 × g and 2 minutes at 750 × g or until flow through was complete. After the addition of 100 μL 0.1% trifluoroacetic acid in HPLC-grade water and centrifugation at 500 × g for 1 min, repeated twice, samples were eluted with 50 μL of 80% acetonitrile/25mM formic acid). Peptides were dried at 45°C for 4 hours in a CentriVap vacuum concentrator (Labconoco). To standardize the peptide loading, the peptide concentration was measured at 480 nm using a Pierce Quantitative Colorimetric Assay (Thermo Fischer Scientific) and Spectra Max M5 (Molecular Devices) and normalized with addition of 3% acetonitrile/0.1% formic acid, to a final concentration of 0.5 μg/μL for CS and IN fractions.

*LC-MS/MS analysis:* Samples were processed with a Dionex UltiMate 3000 RSLC Nano System coupled to the Q Exactive$^{TM}$ HF Hybrid Quadrupole-Orbitrap Mass Spectrometer (ThermoFisher Scientific). 1 μg of peptide was loaded onto a 300μm i.d. × 5mm C18 PepMap 100 trap column. The peptides were washed for 5 minutes flow rate of 5 μL/minute using 98% purified water/2% acetonitrile/0.01% formic acid. The peptides were then processed using a 75 μm × 50 cm reverse-phase Acclaim C18 PepMap 100 analytical column at 50°C. The peptides were separated on the C18 HPLC system using a 120-minute gradient elution method at a flow rate of 300 nl/minute. Mobile phase A was composed of of 0.01% formic acid in water. Mobile phase B comprised of 0.01% formic acid in 80% acetonitrile. Using a linear gradient, the solution was changed from between 2% at time 0, 10% B at 5 minutes, 30% B at 80 minutes, 45% B at 91 minutes, and 100% B at 93 minutes. Between 93 and 98 minutes, the column was held at 100% B. Finally, the solution was changed to 2% B and held for 20 minutes. After HPLC separation, Samples were injected into the Q Exactive HF mass spectrometer through the Nanospray Flex Ion Source fitted with an emission tip (New Objective). With an injection time of 100 ms, the top 20 precursors at 120,000 resolution were monitored.

*LC-MS/MS Analyses:* Mass spectrometry analysis followed.[1] Peaks from the raw file were analyzed by MaxQuant[3,4] (version 1.6.7.0) with settings shown in Table S1 against a FASTA database for *Mus musculus* (downloaded 12/06/2019) with canonical variants and contaminants. Fixed modifications of cysteine carbamidomethylation and variable modifications of oxidation of methionine, hydroxylysine, hydroxyproline, deamidation of asparagine, and conversion of glutamine to pyro-glutamic acid were used.[14] Match-between-runs was used for biological replicates of the same fraction. A decoy database derived from the *Mus musculus* database was used to control the false discovery rate (FDR) to 1%. LFQ was enabled for biological replicates.

**Imaging**.

*Dissection image:* A Leica M80 stereomicroscope (Leica Microsystems) was used to acquire wide-field images of dissected kidneys.

*Immunohistochemistry (IHC):* Cryosections of murine kidneys from E14.5, E18.5, P3, and adult timepoints were acquired at 10 µm thickness using a Shandon Cryotome FE (Thermo), collected on charged slides, and stored at -20°C until processed for IHC.

Incubations were conducted at RT unless indicated otherwise, and samples were protected from light when fluorescent secondary staining reagents were used. After tissues equilibrated to RT, kidney sections were encircled with an ImmEdge pen (Vector Laboratories). The samples were rehydrated in PBS for 10-15 minutes, fixed in 4% paraformaldehyde (PFA; Electron Microscopy Sciences) for 5 minutes, and rinsed in PBS. Sections were blocked for 1 hour with IgG blocking buffer from the Mouse on Mouse (MOM) basic kit (Vector Laboratories) following manufacturer's instructions, rinsed 3 × 2 minutes with PBS, and blocked for 5 minutes with protein diluent from the MOM basic kit. Slides were incubated with primary antibodies (Table S4) in a solution of MOM protein diluent overnight at 4°C, then washed 3 × 2 minutes with 1× PBS. Secondary antibodies, stains, and DAPI (Table S4) were applied to sections in a solution of MOM protein diluent for 45 minutes. Slides were rinsed 3 × 2 minutes with 1 × PBS and tissue was mounted using FluoromountG (Electron Microscopy Sciences) and #1.5 coverslips, and sealed using clear nail polish. Slides were stored at 4°C until imaged with a Leica DMI6000 at 20×

magnification. Negative controls consisted of the same processing with the exclusion of the primary antibodies. Images are representative of ≥ 2 biological replicates for 2 litters.

*3D imaging of decellularized kidneys:* Kidneys were processed for 3D imaging of the ECM using protocols modified from.[5] Depending on the timepoint, dissected kidneys were either directly incubated in SDS solution or embedded in agarose to improve tissue stability (Table S4). The adult timepoint was represented with P28 – P56 kidneys due to more optimum decellularization when the tissues are smaller and as stromal cell accumulations disappear by P28.[6] Low melt agarose (Sigma Aldrich) was suspended in Milli-Q water with 0.02% sodium azide (see concentrations in Table S4), melted and equilibrated to a 37°C water bath and then used to embed the kidney in a cryomold (Tissue-Tek). After equilibrating and solidifying at RT, the agarose embedded samples were transferred to an SDS solution (concentrations, volumes indicated in Table S4). The samples were gently rocked at RT, with daily changes of the SDS solution. Once decellularized, the samples were washed with 1× PBS for 1 hour at RT with rocking. The samples were fixed for 1 hour in 4% PFA in PBS at RT or overnight at 4°C with rocking and washed with 1× PBS for 1 hour RT or 4°C overnight with rocking. Samples were stored in 1× PBS at 4°C until staining. After excess agarose was removed, the samples were transferred to a 96 well plate (volume 150 μL), permeabilized and blocked overnight at 4°C with 10% donkey serum (Lampire Biological Laboratories) and 0.02% sodium azide (Sigma Aldrich) in 0.1% PBST (0.1% Triton-X 100 in 1× PBS). The primary antibodies were diluted to concentrations indicated in Table S4 in 0.2% bovine serum albumin (BSA; Sigma Aldrich) and 0.02% sodium azide in 1× PBS and were added to the sample for 48 hours at 4°C with gentle rocking. After washing for 3 × 30 minutes in 0.1% PBST at RT, the sample was incubated in stains and secondary antibody combinations (Table S4) diluted in 0.2% BSA and 0.02% sodium azide in 1× PBS at 4°C for 48 hours with gentle rocking while protected from light. To provide a general outline of kidney ECM architecture, samples were counterstained with AF488-conjugated wheat germ agglutinin (WGA), which stains a subset of proteoglycans. The samples were washed 3 × 30 minutes in 0.1% PBST at RT and stored in PBS at 4°C with protection from light until imaged.

For imaging, decellularized kidneys were placed in an 8- or 18-well dish (Ibidi) and suspended in 1× PBS at an optimal level that would avoid both dehydration and excessive movement. Kidneys imaged at 40× and 63× were stabilized by bisecting kidneys and wedging the tissue between blocks of 1% agarose. Kidneys were imaged using a Zeiss LSM 880 confocal microscope (Carl Zeiss Microscopy) with Zen 2.3 SP1 black software (V14.0.2.201). Samples were imaged with settings in Table S4. Laser power and gain were optimized for each sample to improve visualization of the ECM. Negative controls consisted of the same process without the addition of the primary antibody and were imaged at the maximum settings for the antibody channels. Images in figures are representative of a minimum of $n$ = 3 biological replicates from a minimum of 2 litters.

*Image processing:* Widefield images and cryosections were processed using FIJI (NIH).[7] Cryosections were processed were optimized per time point to improve visualization (Figure 5) or with the same settings across samples (Figure S5). Confocal z-stacks were processed using nearest-neighbor deconvolution in Zen Blue software (V2.3.64.0; Carl Zeiss Microscopy). In FIJI, decellularized tissues were further processed using bleach correction exponential fit to increase the signal intensity with depth.[8] Kidney z-stacks were visualized using FIJI 3D viewer.[9] Renderings of wide-field images, cryosections, and decellularized tissue were compiled using Adobe Photoshop and Illustrator.

*Statistical analyses:* Analysis of ECM sub-cellular compartments included a two-way ANOVA (factors: fraction and tissue type) with and Tukey or Sidak's multiple comparisons test between tissues with a significance cut-off of $\alpha < 0.05$. ECM functional classifications studied the factor of tissue with an one-way ANOVA and subsequent Tukey multiple comparisons test or unpaired, two tailed t-test with a significance cut-off of $\alpha < 0.05$. To determine if individual proteins in the IN fraction changed over time or with tissue type, one-way ANOVA was performed for data identified in 3-4 timepoints. The data was analyzed using untransformed values if the Brown-Forsythe test for equal standard deviation and the Shapiro Wilk Normality assumptions were met. $Log_2$ transformed data was used if Brown-Forsythe test for equal standard deviation and the Shapiro Wilk normality assumptions were met when the data were transformed. If assumptions were not met, untransformed data was used.[10] Values identified at 2 timepoints were compared using an un-paired two-tailed student t-test using scaled untransformed data if the F-test for variance and Shapiro Wilk test for normality were met. If the assumptions were not met, the values were $log_2$ transformed if the F-test for variance and Shapiro Wilk test for normality were met using the $log_2$ transform data. If both assumptions were not met, untransformed data was used.[10] Significance was determined using a $p$ cut-off of $\alpha < 0.05$. For volcano plot comparisons, Tukey comparisons between all timepoints were performed if identified in 3 or 4 timepoints or un-paired two-tailed student t-test if identified in 2 timepoints. The scaled or $log_2$ transformed data was used based on the assumptions above. Proteins were defined as transiently elevated if a middle timepoint (E18.5 or P3) was significantly higher than an earlier and later timepoint. Imputation was avoided as it negatively affects high abundance proteins in statistical analyses.[11]

**Videos:**

**Video S1: 10× medullary ray sheath fibers at E14.5, E18.5, and P3 stained for COL5, COL4A1, and WGA.** 3D renderings and individual slices of E14.5, E18.5, and P3 kidneys from Figure 7 showed medullary ray sheath fibers. Cortical to the medullary ray sheath fibers were tubules, glomeruli, and the articular artery. 10× confocal z-stacks (x × y) 1.42 mm × 1.42 mm. 3D rendering: z = 100 µm, individual slices: E14.5, z = 576 µm; E18.5 and P3, z = 384 µm**.**

**Video S2: 10× medullary ray sheath fibers at E14.5, E18.5 and P3 stained for COL6, HSPG2, and WGA.** 3D renderings and individual slices of E14.5, E18.5, and P3 kidneys from Figure 7 showed medullary ray sheath fibers. Cortical to the medullary ray sheath fibers were tubules, glomeruli, and the articular artery. 10× confocal z-stacks (x × y) 1.42 mm × 1.42 mm. 3D renderings: z = 100 µm, individual slices: E14.5, z = 307; E18.5 µm and P3, z = 384 µm**.**

**Video S3: 40× medullary ray sheath fibers at E18.5 and P3 stained for COL5, COL4A1, and WGA.** 3D renderings and individual slices of E18.5 and P3 kidneys from Figure 7 showed medullary ray sheath fibers surrounding the developing loop of Henle and collecting duct. 40× insets: 354 × 354 × 45 µm for 3D renderings and individual slices.

**Video S4: 40× vertical fibers in the cortex of E14.5, E18.5, and P3 kidney stained for COL5, COL4A1, and WGA.** 3D renderings of E14.5, E18.5, and P3 kidneys from Figure 8 showed vertical fibers surrounding the nephrogenic zone. 40× insets: 354 × 354 × 45 µm for 3D renderings.

**Tables:**

**Table S1: Raw data supporting the comparison of the matrisome during kidney development.** Includes number of kidneys isolated for mass spectrometry, MaxQuant settings, proteins identified and calculation of cellular compartment and ECM functional classifications, proteins identified for Venn diagram and heat map values, proteins significantly changed or exclusively found at different timepoints, and GO analysis terms.

**Table S2: Raw data supporting the comparison of the matrisome between E14.5 kidneys and whole embryos.** Includes proteins identified and calculation of cellular compartment and ECM functional classifications, proteins significantly changed or exclusively found in whole embryo or kidney, and GO analysis terms.

**Table S3: A common kidney matrisome was observed across different studies.** Comparison of matrisome proteins observed in our study to other kidney matrisome studies showed there was a common set of ECM proteins, and proteins found only during development, in the glomerulus, renal artery, or kidney culture models.

**Table S4: Materials and settings used for imaging the kidney ECM.** Includes optimized tissue processing for decellularization, antibodies and stains used for immunohistochemistry and decellularization, 3D imaging specifications, settings, and locations, and summary of staining patterns observed in immunohistochemistry and decellularization.

**Figures:**

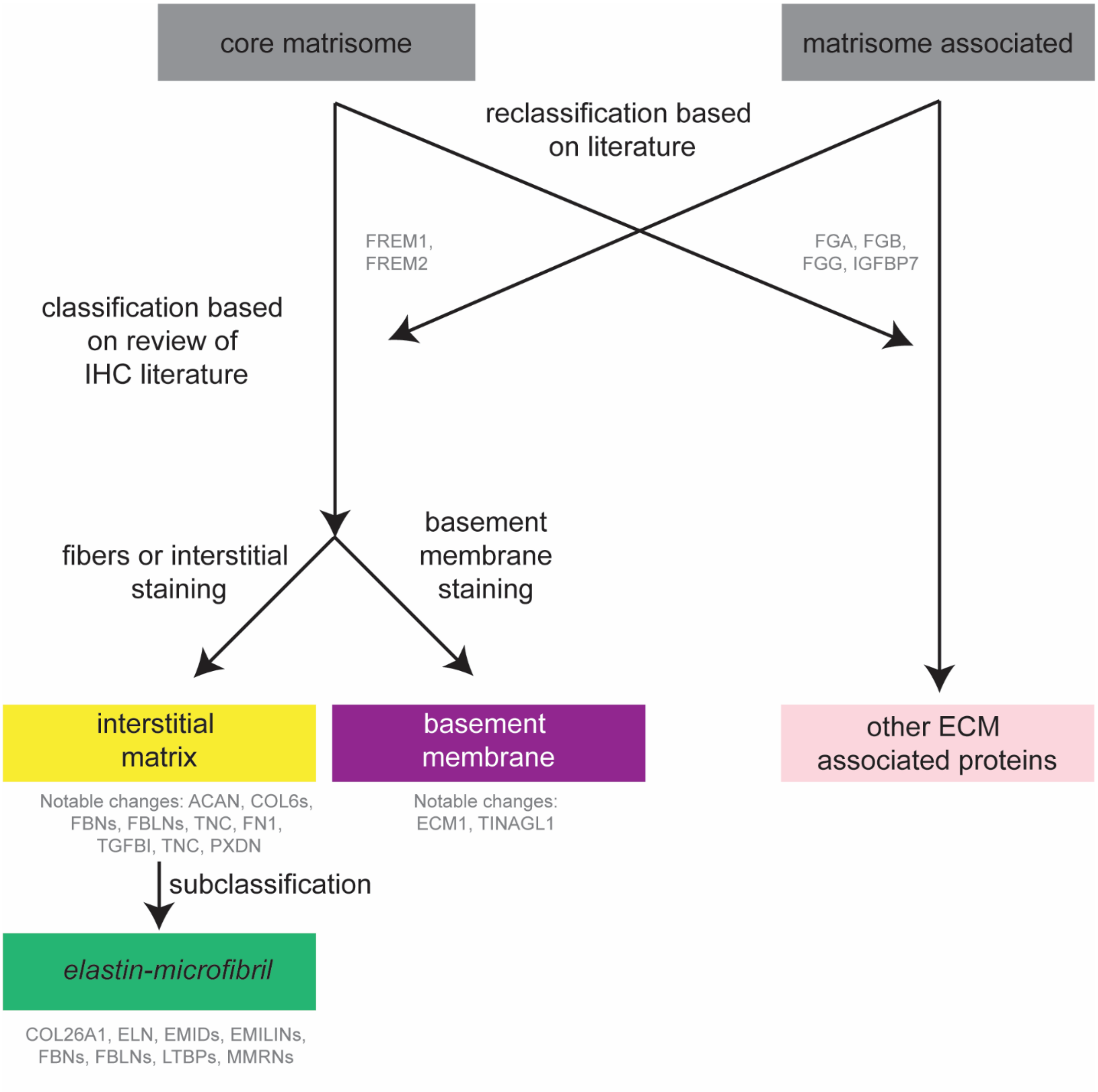


**Figure S1: Flow diagram of the functional classification of proteins as interstitial matrix, basement membrane, and other ECM associated proteins for matrisome identified in this study.** In general, core matrisome proteins were either classified as interstitial matrix or basement membrane, while matrisome-associated proteins were classified as other except for the reclassification of FREM1 and FREM2 as basement membrane proteins. Baseline classifications were modified based on review of the literature.[12-14] If fibers are described (*e.g.* FBN1, TNC, COL6, FN1), it was classified as interstitial matrix. The interstitial matrix was subclassified as *elastin-microfibril axis* based on the literature.[15-27] If described as basement membrane or basement membrane interface, the protein is annotated as basement membrane.

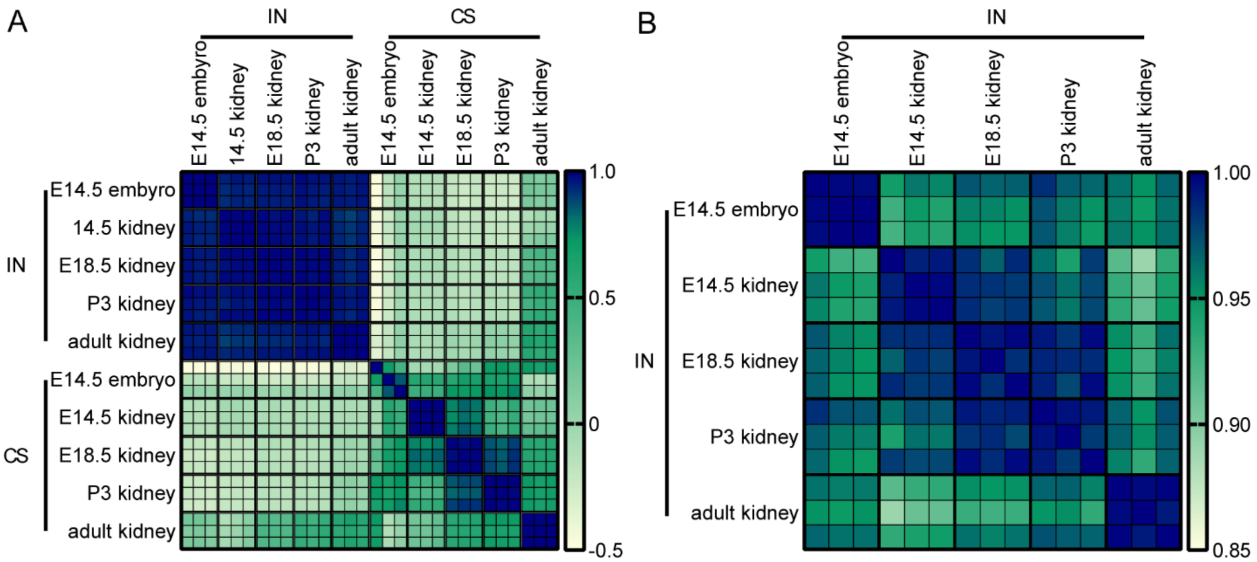


**Figure S2: Proteomic techniques showed high reproducibility between the matrisome for biological replicates. (A)** Comparison of Pearson's correlation coefficients for E14.5, E18.5, P3, adult murine kidneys, and E14.5 whole embryos using scaled LFQ values for matrisome proteins. **(B)** Inset of A focusing on the IN fraction ($n$ = 3 biological replicates per timepoint). Between biological replicates, the Pearson's correlation coefficients were close to 1, indicating the reproducibility of the method, whereas there was greater variation between timepoints.

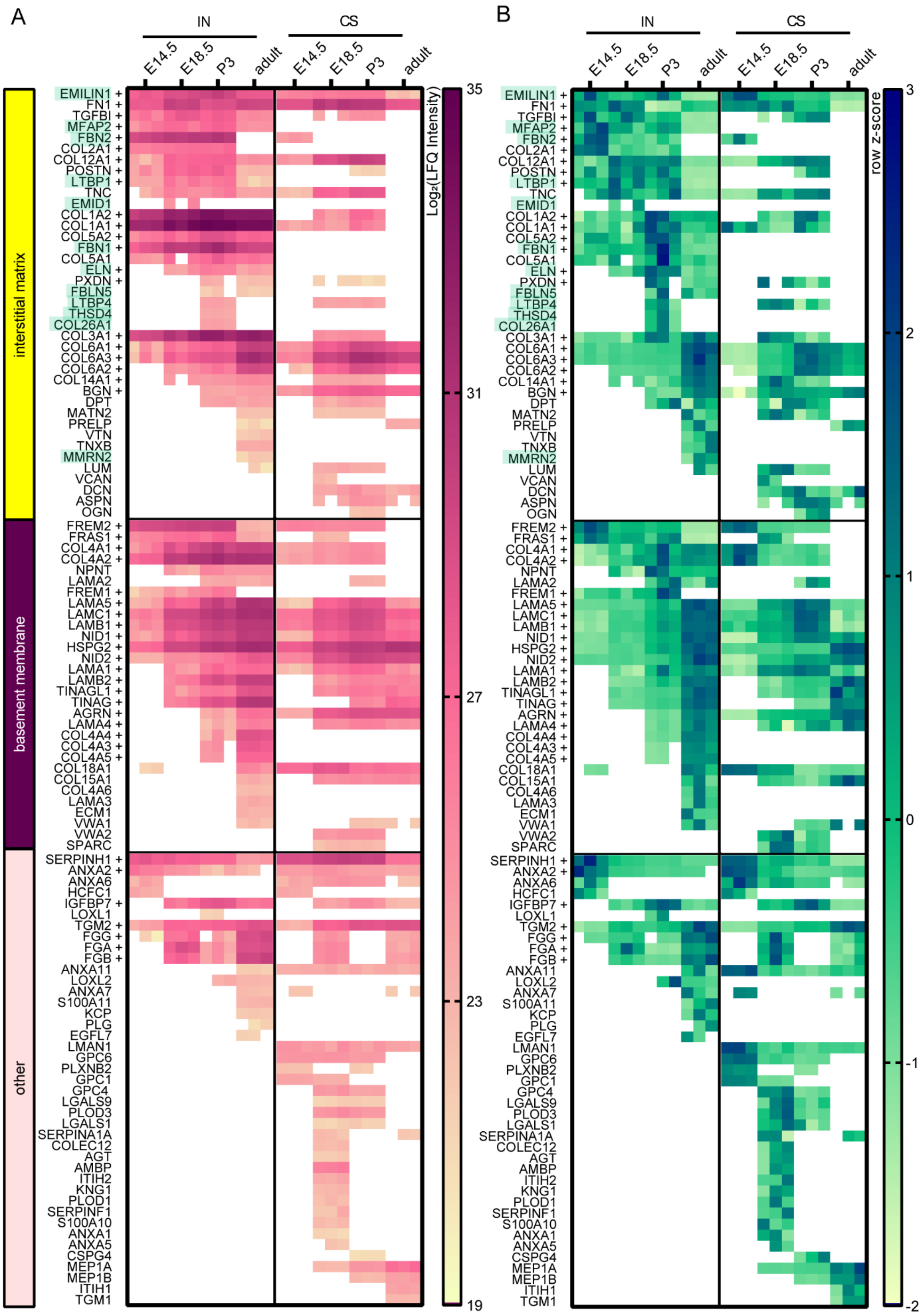

A
IN
CS
E14.5
E18.5
P3
adult
interstitial matrix
basement membrane
other
EMILIN1 +
FN1 +
TGFBI +
MFAP2 +
FBN2 +
COL2A1 +
COL12A1 +
POSTN +
LTBP1 +
TNC
EMID1
COL1A2 +
COL1A1 +
COL5A2 +
FBN1 +
COL5A1
ELN +
PXDN +
FBLN5
LTBP4
THSD4
COL26A1
COL3A1 +
COL6A1 +
COL6A3 +
COL6A2 +
COL14A1 +
BGN +
DPT
MATN2
PRELP
VTN
TNXB
MMRN2
LUM
VCAN
DCN
ASPN
OGN
FREM2 +
FRAS1 +
COL4A1 +
COL4A2 +
NPNT
LAMA2
FREM1 +
LAMA5 +
LAMC1 +
LAMB1 +
NID1 +
HSPG2 +
NID2 +
LAMA1 +
LAMB2 +
TINAGL1 +
TINAG +
AGRN +
LAMA4 +
COL4A4 +
COL4A3 +
COL4A5 +
COL18A1
COL15A1
COL4A6
LAMA3
ECM1
VWA1
VWA2
SPARC
SERPINH1 +
ANXA2 +
ANXA6
HCFC1
IGFBP7 +
LOXL1
TGM2 +
FGG +
FGA +
FGB +
ANXA11
LOXL2
ANXA7
S100A11
KCP
PLG
EGFL7
LMAN1
GPC6
PLXNB2
GPC1
GPC4
LGALS9
PLOD3
LGALS1
SERPINA1A
COLEC12
AGT
AMBP
ITIH2
KNG1
PLOD1
SERPINF1
S100A10
ANXA1
ANXA5
CSPG4
MEP1A
MEP1B
ITIH1
TGM1
Log₂(LFQ Intensity)
35
31
27
23
19
B
IN
CS
E14.5
E18.5
P3
adult
EMILIN1 +
FN1 +
TGFBI +
MFAP2 +
FBN2 +
COL2A1 +
COL12A1 +
POSTN +
LTBP1 +
TNC
EMID1
COL1A2 +
COL1A1 +
COL5A2 +
FBN1 +
COL5A1
ELN +
PXDN +
FBLN5
LTBP4
THSD4
COL26A1
COL3A1 +
COL6A1 +
COL6A3 +
COL6A2 +
COL14A1 +
BGN +
DPT
MATN2
PRELP
VTN
TNXB
MMRN2
LUM
VCAN
DCN
ASPN
OGN
FREM2 +
FRAS1 +
COL4A1 +
COL4A2 +
NPNT
LAMA2
FREM1 +
LAMA5 +
LAMC1 +
LAMB1 +
NID1 +
HSPG2 +
NID2 +
LAMA1 +
LAMB2 +
TINAGL1 +
TINAG +
AGRN +
LAMA4 +
COL4A4 +
COL4A3 +
COL4A5 +
COL18A1
COL15A1
COL4A6
LAMA3
ECM1
VWA1
VWA2
SPARC
SERPINH1 +
ANXA2 +
ANXA6
HCFC1
IGFBP7 +
LOXL1
TGM2 +
FGG +
FGA +
FGB +
ANXA11
LOXL2
ANXA7
S100A11
KCP
PLG
EGFL7
LMAN1
GPC6
PLXNB2
GPC1
GPC4
LGALS9
PLOD3
LGALS1
SERPINA1A
COLEC12
AGT
AMBP
ITIH2
KNG1
PLOD1
SERPINF1
S100A10
ANXA1
ANXA5
CSPG4
MEP1A
MEP1B
ITIH1
TGM1
row z-score
3
2
1
0
-1
-2

**Figure S3: Heat map comparison of the dynamic changes of the matrisome for both the IN and CS fraction**. **(A)** The raw intensity of ECM proteins was higher in the IN compared to the CS fraction. **(B)** Row z-score heatmap comparison of IN and CS fractions, based on scaled LFQ intensity, showed similar trends in protein expression. Heat maps were manually clustered based on classification as interstitial matrix, basement membrane, or other ECM associated proteins. *Elastin-microfibril axis* proteins are highlighted in green. + indicates $p < 0.05$ for proteins identified in the IN fraction based on one-way ANOVA (identified in 3 - 4 timepoints) or unpaired, two tailed t-test (identified in 2 timepoints). Proteins identified in $n \geq 2$ biological replicates were included in the heat map analysis. White boxes signify zero intensity values.

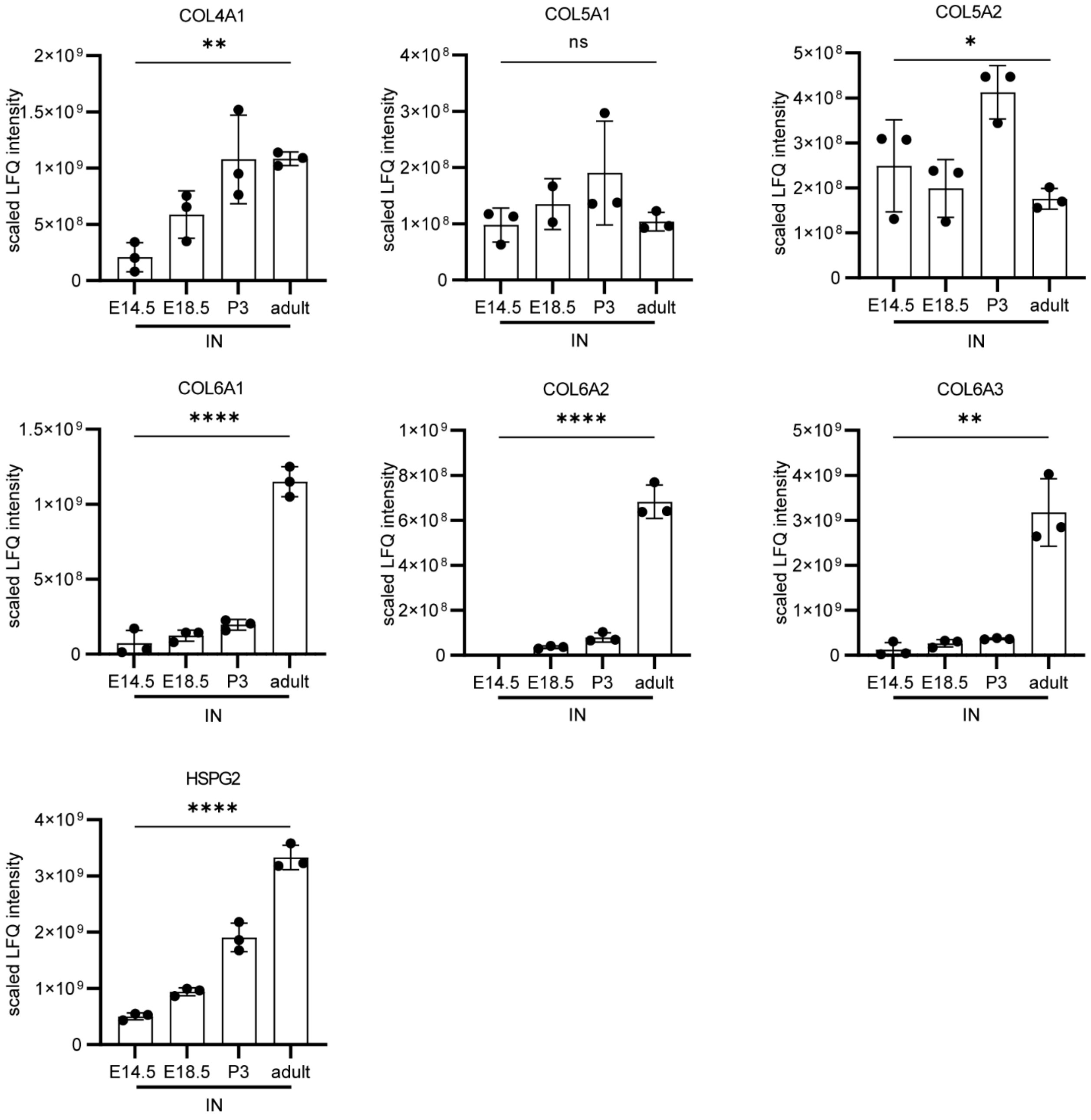


**Figure S4: Scaled LFQ intensity values for the IN fraction for proteins of interest visualized using IHC.** Significance, determined by one-way ANOVA, is indicated on the graph as follows: $p < 0.05$ = *, $p < 0.01$ = **, $p < 0.001$ = ***, $p < 0.0001$ = ***, $p < 0.00001$ = ****, and non-significant = ns. Tukey comparisons between timepoints are found in Table S1.

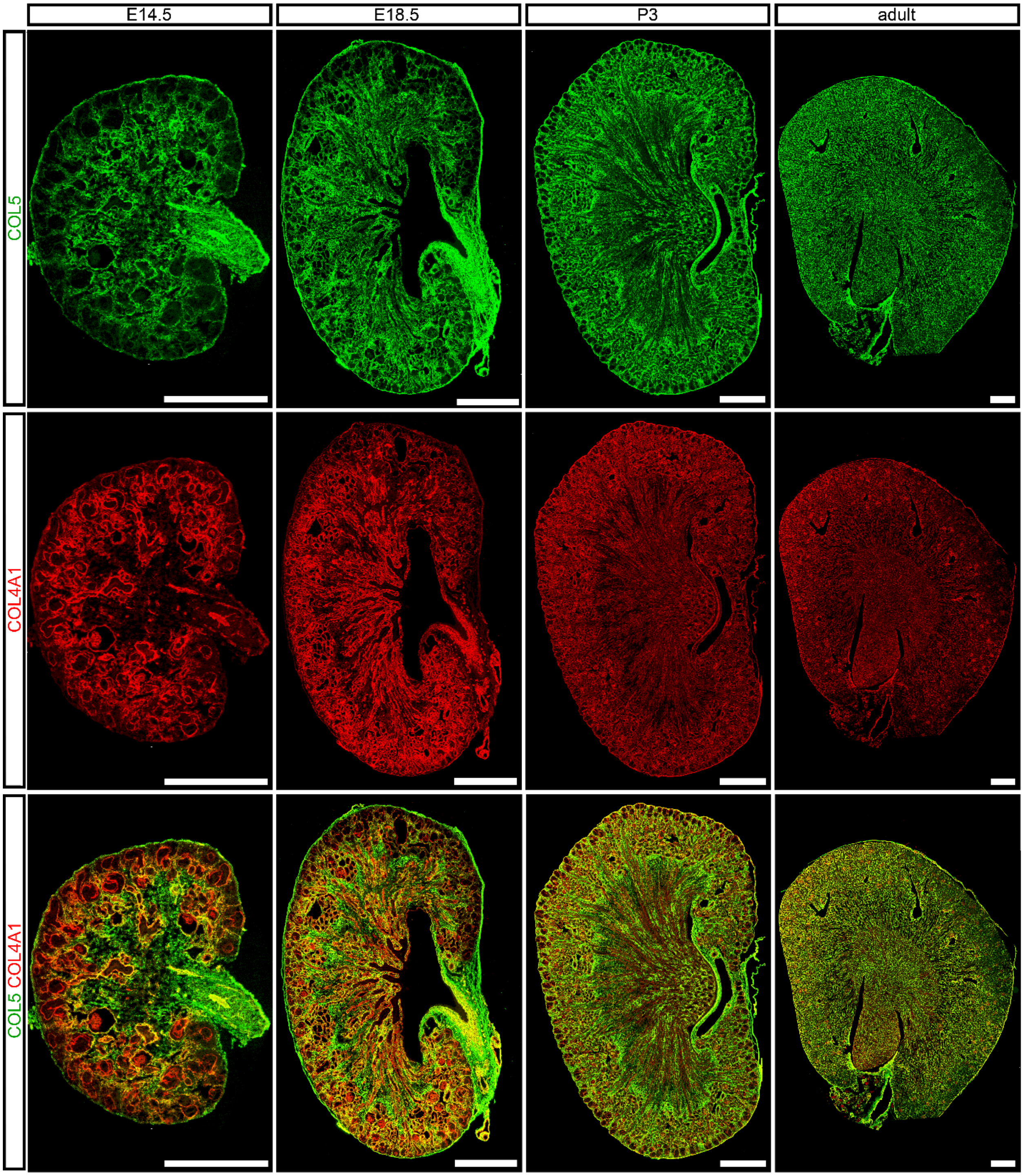


**Figure S5: Spatiotemporal change of COL5, COL4A1 in murine developing kidney.** Cryosections of E14.5 – adult kidneys showed a decrease interstitial matrix space (green = COL5)

and a corresponding increase in basement membrane area (red = COL4A1;). Scale bars = 500 µm. Representative images from *n* = 2 biological replicates.

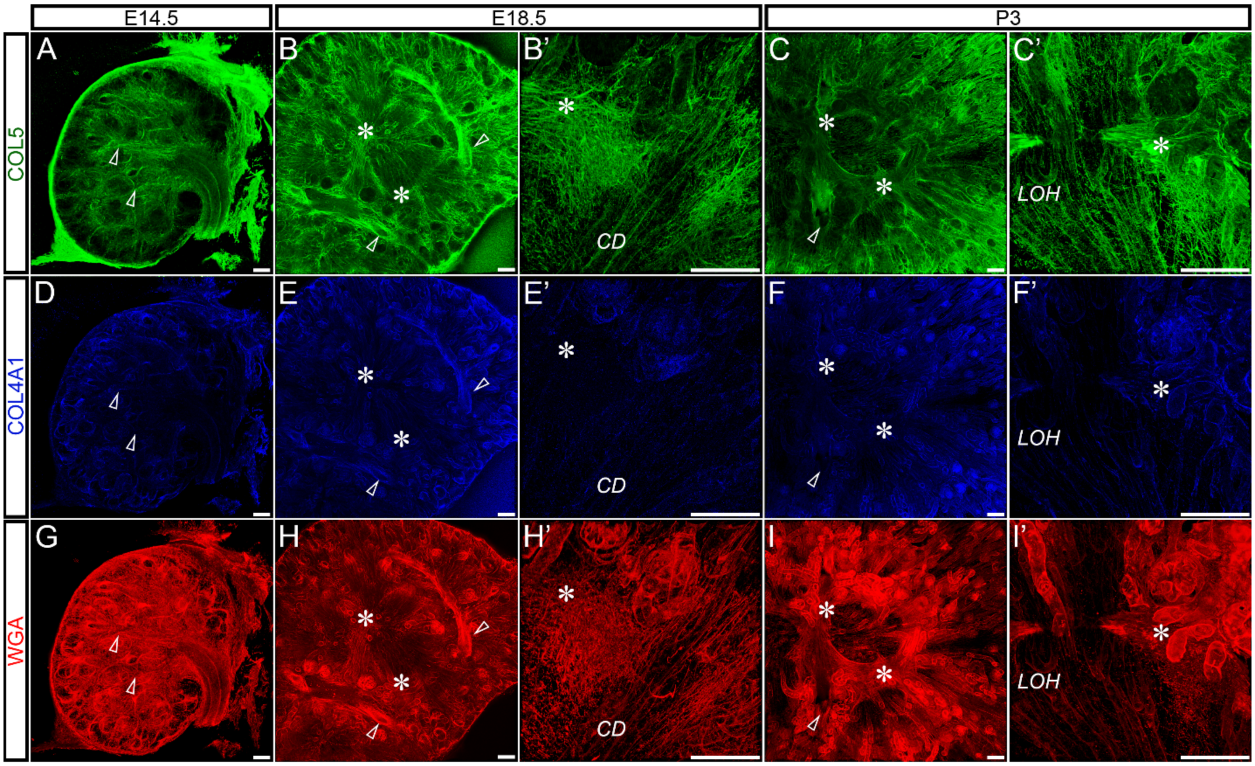


**Figure S6: Medullary ray sheath fibers were only found at E18.5 and P3 and stained for COL5**. **(A, D, G)** At E14.5, a fibrous ECM (green = COL5) ran parallel to developing blood vessels and surrounding tubules (blue = COL4A1, red = WGA). **(B, B', C, C', E, E', F, F', H, H', I, I')** Medullary ray sheath fibers (*) surrounded the developing nephron (*CD* = collecting duct, *LOH* = loop of Henle) at E18.5 – P3. Scale bars = 100 µm. Dimensions of 10× confocal z-stacks in A-I: 1.42 mm × 1.42 mm × 100 µm ($x \times y \times z$); in B', C', E', F', H', and I', 40× insets: 354 × 354 × 45 µm. Open arrowheads = blood vessels. Representative images from *n* = 3 biological replicates.

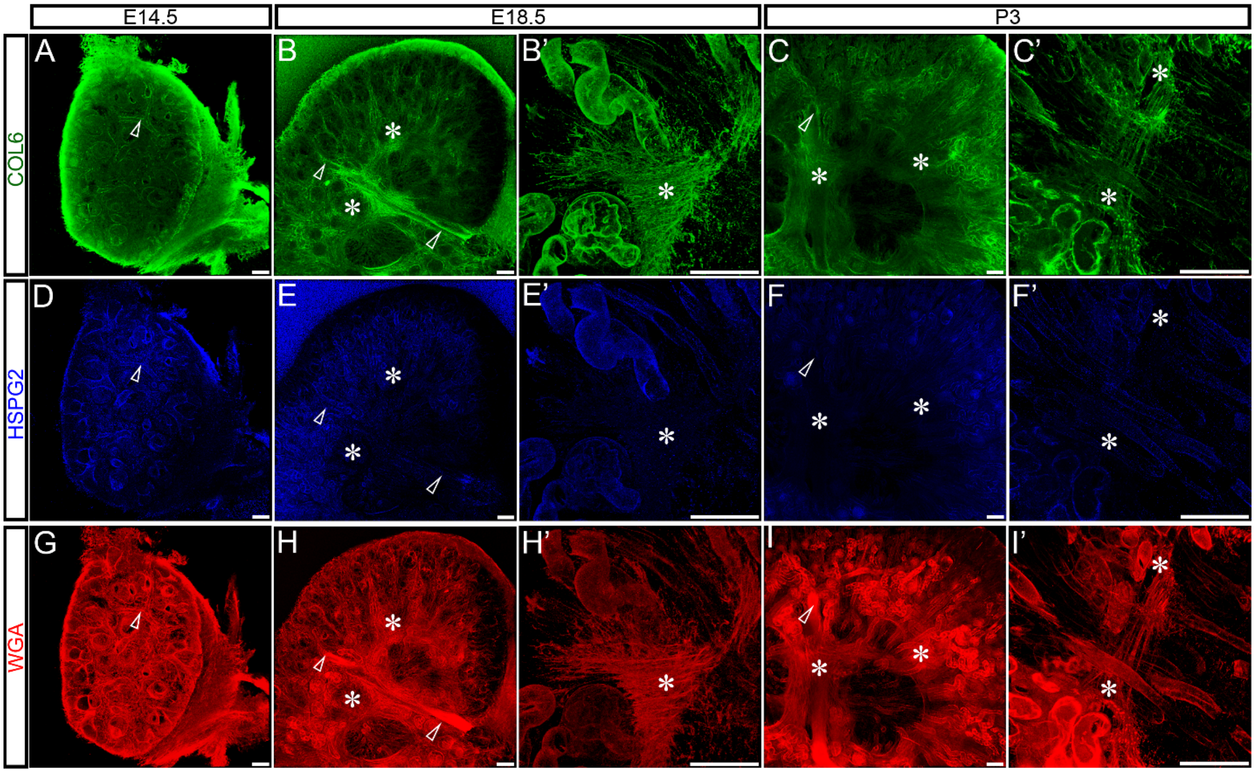


**Figure S7: Medullary ray sheath fibers were only found at E18.5 and P3 and stained for COL6**. **(A, D, G)** At E14.5, a fibrous ECM (green = COL6) ran parallel to developing blood vessels and surrounding tubules (blue = HSPG2; red = WGA). **(B, B’, C, C’, E, E’, F, F’, H, H’, I, I’)** Medullary ray sheath fibers (*) surrounded the developing nephron (*CD* = collecting duct, *LOH* = loop of Henle) at E18.5 – P3. Scale bars =100 µm. Dimensions of 10× confocal z-stacks in A-I: 1.42 mm × 1.42 mm × 100 µm (*x* × *y* × *z*); in B’, C’, E’, F’, H’, and I’, 40× insets: 354 × 354 × 36 µm. Open arrowheads = blood vessels. Representative images from $n = 3$ biological replicates.

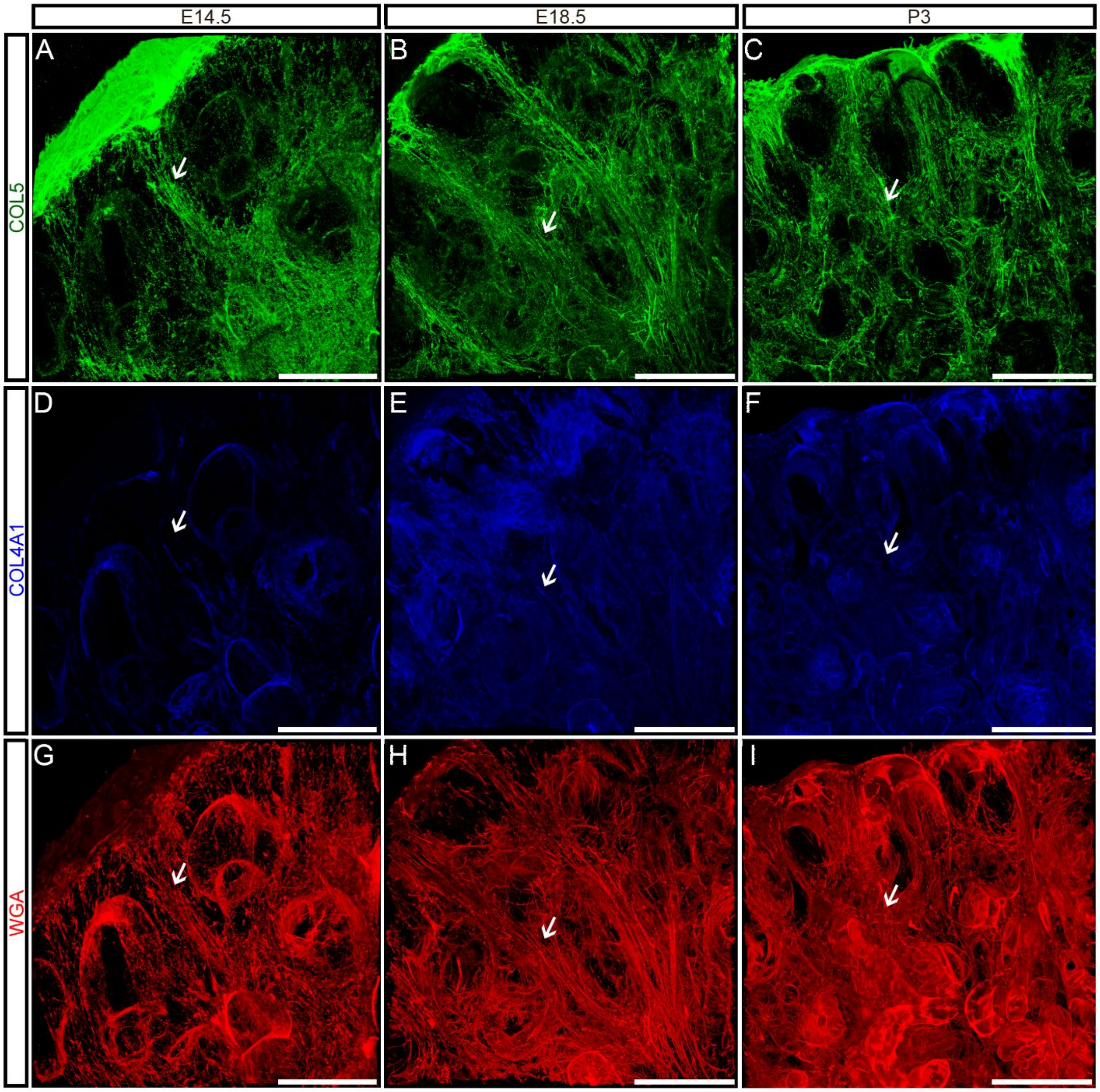


**Figure S8: COL5 vertical fibers were observed in the cortex.** E14.5, E18.5, and P3 vertical fibers (arrows) ran parallel to, and emanated from, the tubules to the capsule in vertical fibers (green = COL5; blue = COL4A1; red = WGA). Dimensions of 40× confocal z-stacks 354 × 354 × 45 µm (*x* × *y* × *z*). Representative image from *n* = 3 biological replicates.

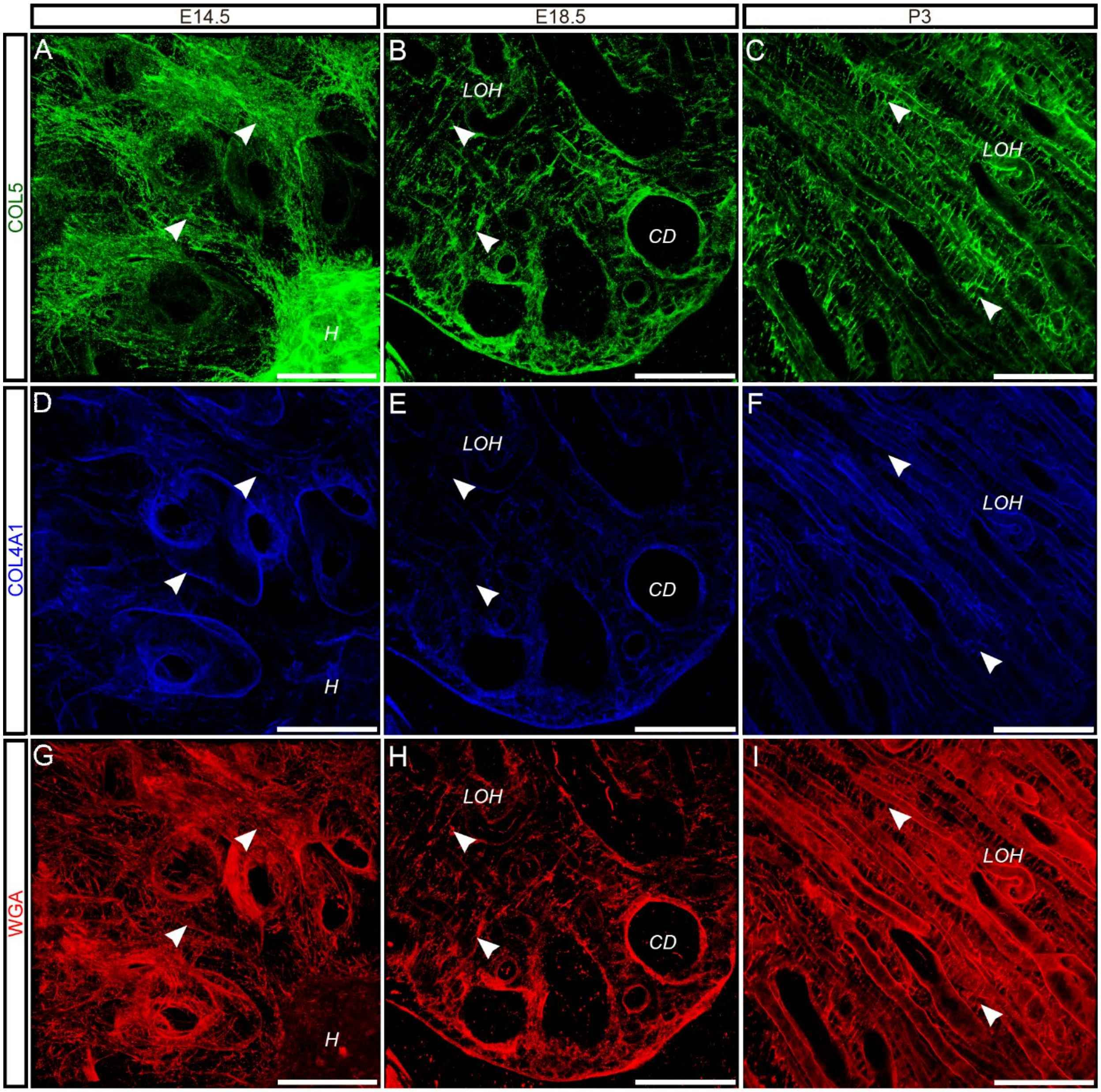


**Figure S9: COL5 "rungs of a ladder" fibers were observed in the medulla.** (**A, B, G**) Some E14.5 fibers (arrows) extended from the developing renal hilum (*H*). **(B, C, E, F, H, I) "**Rungs of a ladder" appearance of the interstitial matrix (arrowheads) at E18.5 and P3 between loop of Henle (*LOH*) and collecting duct (*CD*) tubules. Scale bars = 100 µm. Dimensions of 40× confocal z-stacks in A, D, G: 354 × 354 × 45 µm ($x \times y \times z$); B, C, E, F, H, I: 354 × 354 × 11 µm. Representative image from $n$ = 3 biological replicates.

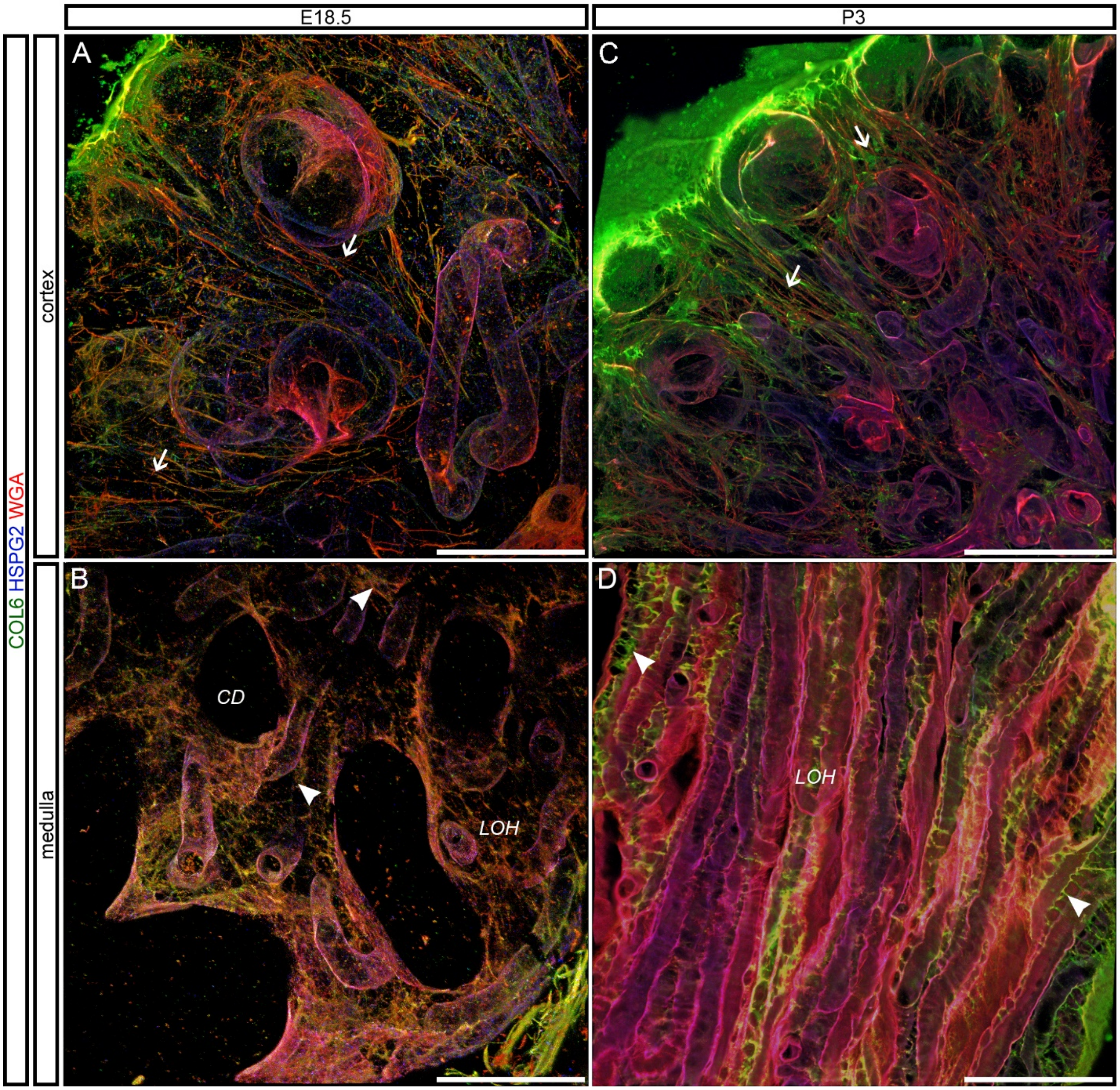


**Figure S10: The morphology of interstitial fibers was distinct between cortex and medulla for COL6. (A, C)** E18.5, and P3 vertical fibers (arrows) ran parallel to and emanated from the tubules to the capsule in vertical fibers (green = COL6; blue = HSPG2; red = WGA). **(B-D) "**Rungs of a ladder" appearance of the interstitial matrix (arrowheads) at E18.5 and P3 between loop of Henle (*LOH*) and collecting duct (*CD*) tubules. Scale bars = 100 μm. Dimensions of 40× confocal z-stacks in A, C: 354 × 354 × 45 µm ($x \times y \times z$); B, D: 354 × 354 × 11 µm. Representative image from $n = 3$ biological replicates.

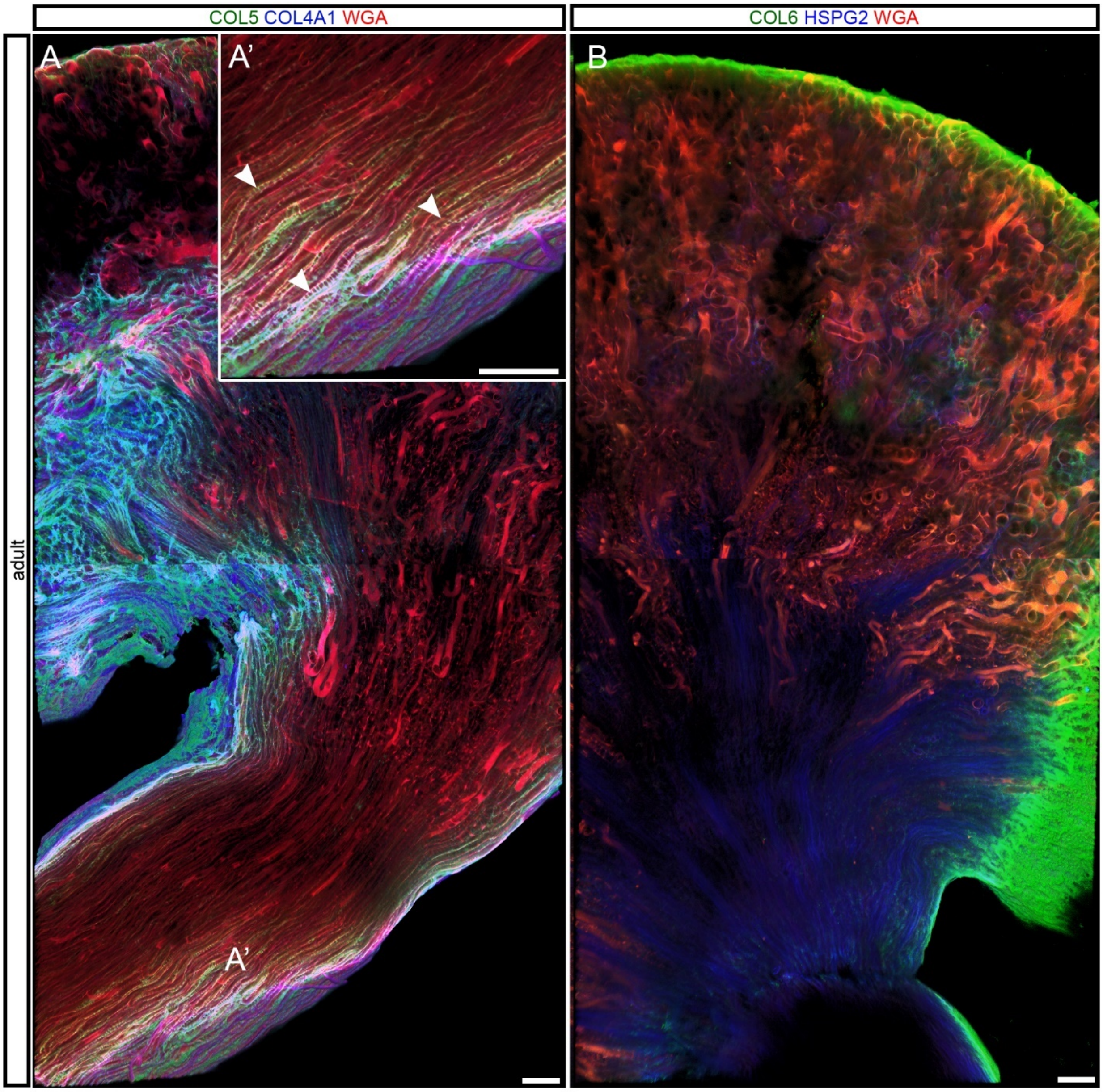


**Figure S11: Interstitial accumulations were not observed in the adult mouse kidney. (A, B)** Adult kidneys stained for COL5, COL4A1 and WGA (A) and COL6, HSPG2, and WGA (B) do not show accumulations of interstitial matrix as observed at developmental timepoints. **(A')** Inset showed "rungs of a ladder" fibers (arrowhead) were observed in the papilla**.** (A, B) 10× confocal z-stacks 2.8 mm × 1.4 mm × 100 µm ($x \times y \times z$). Representative image from $n = 3$ biological replicates.
